# Reelin marks cocaine-activated striatal ensembles, promotes neuronal excitability, and regulates cocaine reward

**DOI:** 10.1101/2024.06.17.599348

**Authors:** Kasey L. Brida, Emily T. Jorgensen, Robert A. Phillips, Catherine E. Newman, Jennifer J. Tuscher, Emily K. Morring, Morgan E. Zipperly, Lara Ianov, Kelsey D. Montgomery, Madhavi Tippani, Thomas M. Hyde, Kristen R. Maynard, Keri Martinowich, Jeremy J. Day

## Abstract

Drugs of abuse activate defined neuronal ensembles in brain reward structures such as the nucleus accumbens (NAc), which are thought to promote the enduring synaptic, circuit, and behavioral consequences of drug exposure. While the molecular and cellular effects arising from experience with drugs like cocaine are increasingly well understood, the mechanisms that sculpt NAc ensemble participation are largely unknown. Here, we leveraged unbiased single-nucleus transcriptional profiling to identify expression of the secreted glycoprotein Reelin (encoded by the *Reln* gene) as a marker of cocaine-activated neuronal ensembles within the rat NAc. Multiplexed in situ detection confirmed selective expression of the immediate early gene *Fos* in *Reln+* neurons after cocaine experience, and also revealed enrichment of *Reln* mRNA in *Drd1*+ medium spiny neurons (MSNs) in both the rat and human brain. Using a novel CRISPR interference strategy enabling selective *Reln* knockdown in the adult NAc, we observed altered expression of genes linked to calcium signaling, emergence of a transcriptional trajectory consistent with loss of cocaine sensitivity, and a striking decrease in MSN intrinsic excitability. At the behavioral level, loss of *Reln* prevented cocaine locomotor sensitization, abolished cocaine place preference memory, and decreased cocaine self-administration behavior. Together, these results identify Reelin as a critical mechanistic link between ensemble participation and cocaine-induced behavioral adaptations.

## INTRODUCTION

Reward-linked adaptive and learned behaviors are regulated by mesolimbic dopamine circuits in the brain^1–4^. These circuits consist of dopaminergic neurons in the ventral tegmental area (VTA) that densely innervate the nucleus accumbens (NAc), a subregion of the striatum that directly shapes motivated behaviors^5^. Mesolimbic dopamine circuitry is a crucial target for many drugs of abuse^3^, which act through distinct transporters, ion channels, and receptors to increase dopamine neurotransmission in the NAc^6,7^. On short timescales, this increase in dopaminergic signaling promotes the excitability of medium spiny neurons (MSNs) in the NAc^8^, leading to calcium influx and altered activity patterns^9–11^. Over longer timescales (minutes to hours), dopamine receptor activation initiates well-defined signal transduction cascades that converge to regulate expression of immediate early gene (IEG) programs strongly linked to synaptic and behavioral plasticity^12–16^. Together, these mechanisms serve as fundamental and conserved biochemical steps that link drug experience to the enduring molecular and physiological changes found in substance use disorders^17^.

Despite the robust elevation in NAc dopamine produced by psychostimulant drugs like cocaine, emerging evidence suggests that cocaine experience activates a small proportion (∼10-20%) of N Acneurons, comprising cocaine-sensitive ensembles^9,10,18–20^. Though small in number, these ensembles exhibit strong control over drug-related behaviors. Selective inactivation of neurons engaged by cocaine abolishes subsequent sensitized behavioral responses^21–23,24^, and modulates cocaine seeking^25^. While neuronal subtypes that participate in drug-responsive ensembles varies based on drug class^10^, cocaine preferentially affects dopamine receptor type-1 class medium spiny neurons (D1-MSNs^9,26^), leading to prolonged increases in excitability^8,27,28^, transcription of IEGs^29–31^, and altered synaptic plasticity^32–34^ that drives prolonged behavioral adaptations produced by drug exposure^35–37^. However, even within the D1-MSN subpopulation, response to cocaine is highly heterogeneous^9^, and the molecular mechanisms that regulate drug ensemble participation in the NAc remain unknown.

Here, we leveraged unbiased single-nucleus RNA sequencing (snRNAseq) datasets collected after cocaine exposure^20,38^ to identify markers of cocaine-activated D1-MSNs in the NAc. Surprisingly, mRNA for the secreted extracellular matrix protein Reelin (encoded by the *Reln* gene) was enriched in D1-MSNs that were activated by cocaine. CRISPR-based knockdown of *Reln* decreased transcriptional states associated with activation by cocaine, altered expression of ion channels related to neuronal excitability, and impaired MSN excitability. Moreover, loss of *Reln* in the NAc abolished locomotor sensitization and place preference for cocaine, and dampened cocaine self-administration behavior. Together, these results identify *Reln* as a stable marker of cocaine ensembles and reveal a key role for *Reln* in the transcriptional, electrophysiological, and behavioral properties of cocaine-induced striatal plasticity.

## RESULTS

### Reln expression marks a cocaine-sensitive population of neurons in the NAc

Although cocaine promotes robust increases in dopamine capable of activating low-affinity DRD1 dopamine receptors on D1-MSNs, only a small fraction of D1-MSNs respond to cocaine as defined by increased calcium influx or IEG expression^9,39^. This small population of activated neurons constitutes an ensemble that critically contributes to synaptic, cellular, and behavioral adaptations produced by cocaine^9,21,26^. Despite the central role of this ensemble, prior studies have not identified potential markers for this labile population that would predict ensemble membership. Previously, we identified a subpopulation of D1-MSNs within the NAc that exhibits a robust transcriptional response following acute and repeated cocaine exposure using snRNA-seq^20,38^. Using these published datasets, we took an unbiased approach to identify a marker gene for the subset of cocaine-sensitive cells within this D1-MSN subtype (**Fig. 1a,b; Fig. S1a**). We first subclustered all D1-MSNs to isolate the activated subpopulation from inactive populations (i1-i4; **Fig. 1b**). Activation was based on the average expression of significant cocaine differentially expressed genes (DEGs; **Fig. 1g**). Assessing differential transcript enrichment within the activated cluster, we identified *Reln* as a potential marker of this activated population (**Fig. 1c-f**). Over 80% of activated D1-MSNs expressed *Reln* mRNA, with average expression ∼10 times higher compared to inactive clusters (**Fig. 1d-f**). *Reln* expression was also significantly correlated with composite IEG expression (**Fig. 1f**). When stratifying all D1-MSNs based on *Reln* expression, only *Reln*+ D1-MSNs demonstrated robust induction of IEGs, such as *Fos* (**Fig 1g,h; Fig. S1**). *Reln* levels were not altered by cocaine administration (**Fig. 1h**). These results were consistent when data was partitioned by acute or repeated exposure, though some IEGs exhibited exposure-dependent induction properties (**Fig. S2**). Notably, cocaine-responsive D1-MSNs did not exhibit enrichment for *Creb1* mRNA (**Fig. 1d**), which encodes an activity-dependent transcription factor (CREB) previously found to regulate memory “engram’ participation in other brain regions^40,41,42^. Similarly, activation by cocaine was not predicted by expression of the *Drd1* dopamine receptor itself, which was consistently abundant across D1-MSN subclusters (**Fig. 1d**). While D1-MSNs are the principal neuronal population activated by cocaine, a small percentage of D2-MSNs also show increased IEGs. *Reln* similarly marks these cocaine-sensitive D2-MSNs, suggesting it may serve as a broad marker of cocaine-sensitivity (**Fig. S3**).

**Figure 1.**
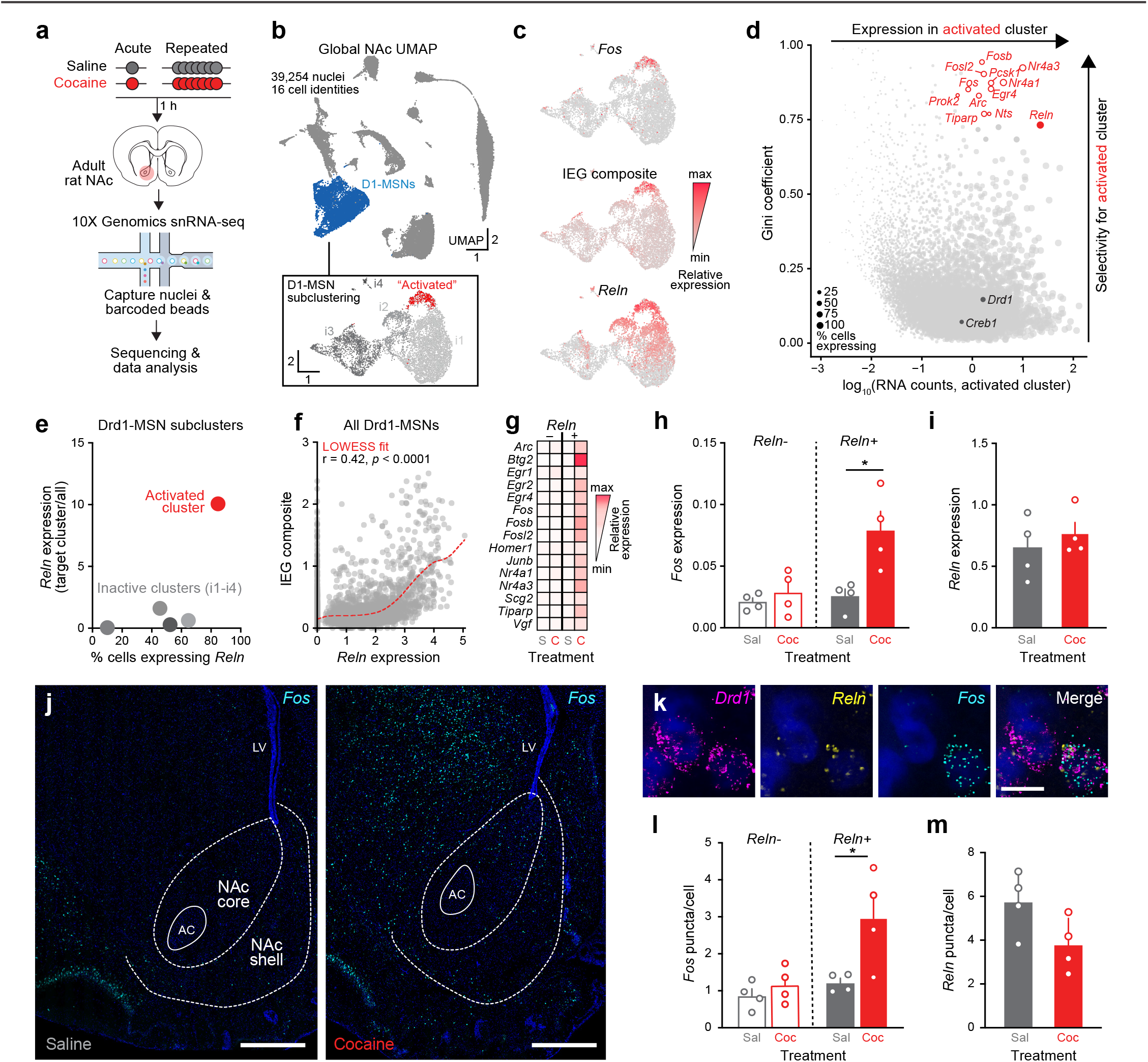
*Reln* marks cocaine-sensitive *Drd1+* MSNs. **a**, snRNA-seq workflow for acute and repeated saline or cocaine. **b**, Top: Uniform Manifold Projection Approximation (UMAP) of integrated NAc snRNA-seq. Bottom: Subclustered D1-MSNs (active = a1; inactive = i1-4). **c**, Feature plot showing concentration of mRNA for *Fos* and other IEGs (as a composite) in the a1 population, which also expresses *Reln* mRNA at high levels. **d**, Gini coefficient analysis comparing D1-MSN subclusters identifies *Reln* mRNA as both a highly selective and highly abundant marker of the activated subcluster. Cocaine-responsive imediate early genes are shown in open circles. **e**, Scatter plot showing the percent of cells within a given cluster expressing *Reln* by the relative expression of *Reln* within a given cluster as compared to all other clusters. The a1 cluster has the highest percentage of cells expressing *Reln* and demonstrates the highest levels of *Reln* expression. **f**, Across D1-MSNs, *Reln* mRNA and composite IEG expression are positively correlated, with cells expressing the highest level of *Reln* exhibiting the strongest transcriptional change to cocaine. Dashed red line shows LOWESS fit. **g**, Heatmap of IEG expression split by treatment and *Reln* expression status. Data are mean values from all Drd1-MSNs, normalized to row average. **h**, Quantification of *Fos* in D1-MSNs, split by *Reln* expression and treatment. snRNA-seq data show *Reln*+ D1-MSNs have a robust transcriptional response to cocaine, while *Reln*-cells do not (p = 0.015, nested one-way ANOVA with Tukey’s multiple comparisons correction). **i**, *Reln* expression does not differ between saline and cocaine-treated animals (nested t-test, p = 0.49). **j**, smRNA-FISH probing for *Fos* in saline animal (right) and cocaine animal (left; scale bar, 1 mm). **k**, High-magnification images from the NAc core of multiplexed smRNA-FISH probing for *Drd1, Reln*, and *Fos* in cocaine-treated animal (scale bar, 10 µm). **l**, Quantification of *Fos* puncta in *Drd1*+ cells, split by *Reln* expression. *Reln*+ cells have a significant transcriptional response to cocaine, while *Reln*-cells do not (p = 0.0433, nested one-way ANOVA with Tukey’s multiple comparisons correction). **m**, *Reln* expression in *Drd1*+ cells in the NAc does not change following cocaine injection (p = 0.08, nested t-test).

As cocaine-linked adaptations primarily depend on D1-MSNs, we focused on this population for in situ validation of *Reln* as a marker of cocaine-sensitivity using single-molecule RNA-FISH (smRNA-FISH). As in the acute snRNA-seq experiment, animals received a single intraperitoneal (i.p.) injection of either saline or 20 mg/kg cocaine 1 hour prior to tissue collection. We then performed smRNA-FISH, probing for *Drd1, Reln*, and *Fos* (**Fig. 1j**) expression. Analysis was restricted to the NAc for direct comparison to snRNA-seq data. We observed a significant increase in *Fos*+ cells in the cocaine group compared to the saline group (**Fig. S1k**). As in the snRNA-sequencing data, *Reln*+/*Drd1+* cells have a significant increase in *Fos* expression compared to saline control, while *Reln*-/ *Drd1+* cells lack this response (**Fig. 1l**). Similarly, *Reln* levels remained unchanged in D1-MSNs from rats treated with cocaine (**Fig. 1m**).

A parsimonious explanation for *Reln* enrichment in cocaine-activated MSNs is that *Reln* is either an activity-responsive IEG itself, or is a direct target of transcription factors that are rapidly activated by cocaine. Notably, snRNAseq (**Fig. 1i**) and smRNA-FISH (**Fig. 1m**) data both demonstrate that *Reln* expression is unaltered in D1-MSNs following cocaine experience, at the same timepoint where other IEGs are reliably detected. Additionally, re-analysis of prior work^20^ revealed that *Reln* mRNA is not altered in primary striatal neurons following direct dopamine exposure (**Fig. S4**), further suggesting that *Reln* is not an IEG. To investigate whether *Reln* may be a target of core transcription factors induced by cocaine and dopamine, we analyzed previously published datasets using a multiplexed CRISPR activation strategy to simultaneously overexpress key dopamine-induced transcription factors^20^, including AP-1 factors (*Fos, Fosb*, and *Junb*), Early Growth Response factors (*Egr2, Egr3*, and *Egr4*), and Nuclear Receptor subfamily 4 group A factors (*Nr4a1* and *Nr4a2*). Critically, *Reln* mRNA expression as detected by RNA-seq remained unchanged following this manipulation (**Fig. S4**). Together, these observations support the interpretation that *Reln* is a stable marker of D1-MSNs that are capable of participating in drug ensembles rather than a cocaine-responsive transcript.

### Reln is enriched in D1-MSNs in both the rat and human brain, and exhibits graded expression across the striatum

Although Reelin’s roles in the cortex and hippocampus have been thoroughly studied^43–46^, much less is known about its cellular and spatial distribution in the striatum. To further characterize the localization of *Reln* mRNA, we performed smRNA-FISH from rat coronal brain sections, probing for *Drd1, Drd2*, and *Reln*. Throughout the striatum, more *Drd1*+ cells were also *Reln*+ as compared to *Drd2*+ cells, and *Drd1*+ cells also expressed more *Reln* transcripts per cell (**Fig. S5b**). This was also true of the NAc (**Fig. 2a,b**). The striatum can be divided into anatomical subregions with different contributions to behavioral output^47,48^. Therefore, we assessed *Reln* expression across striatal subregions (NAc core, NAc shell, ventrolateral striatum (VLS), ventromedial striatum (VMS), dorsolateral striatum (DLS), and dorsomedial striatum (DMS)). This comparison revealed a gradient of *Reln+* cells, with a higher percent of *Reln*+ cells in the dorsal striatum and nearly absent *Reln* expression in the NAc shell (**Fig. S5c-g**). This dorsal-ventral gradient of *Reln* expression is observed in both *Drd1* and *Drd2* neurons, though *Drd1+/Reln*+ cells outnumber *Drd2+/Reln*+ cells in every subregion except the shell (**Fig. S5c**). As in the NAc as a whole, *Drd1+* cells expressed more *Reln* compared to *Drd2+* cells in the dorsal striatum (**Fig. S5d**). *Reln* expression also exhibits lateralization, with more *Reln+* positive cells located in lateral subregions of the striatum as compared to medial subregions (**Fig. S5e**). No sex differences in *Reln* expression were detected at either the anatomical or cellular level (**Fig. S5h-k**).

**Figure 2.**
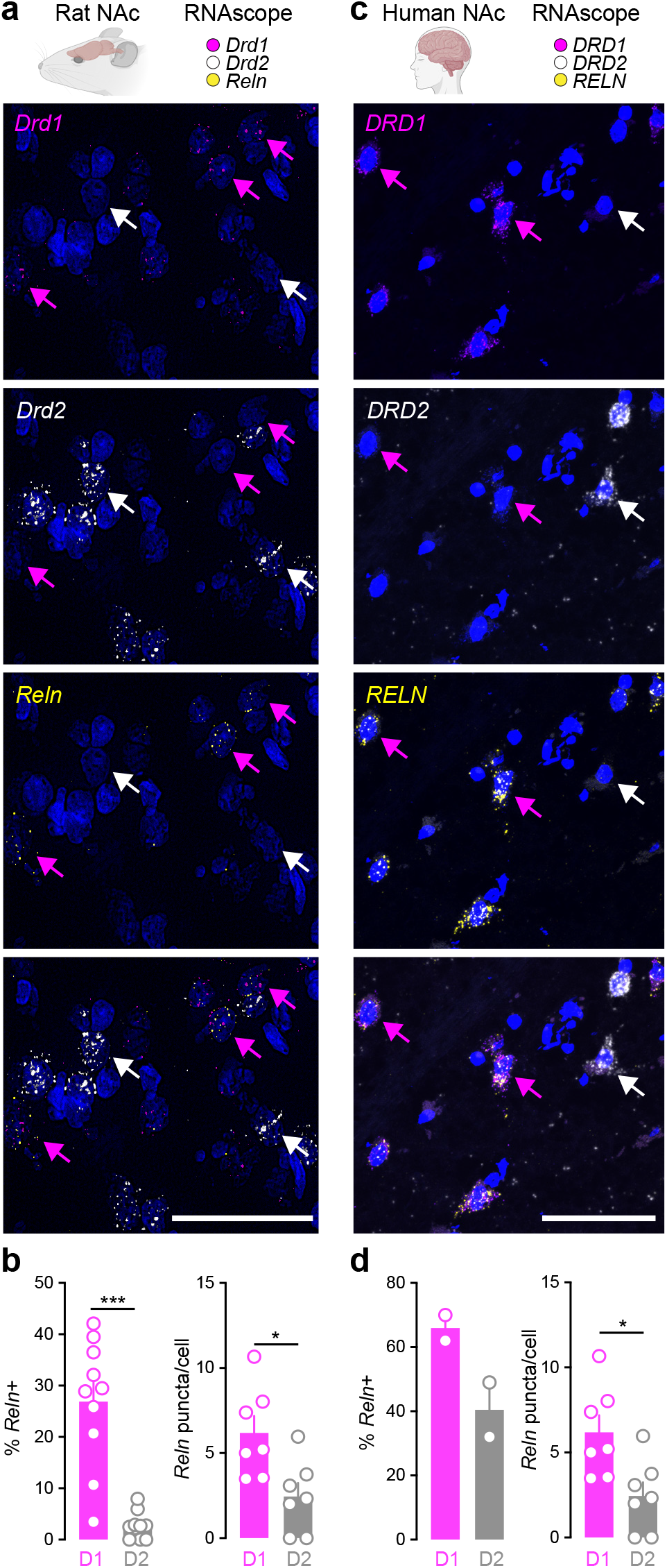
Conserved enrichment of *Reln* mRNA in *Drd1*-MSNs in both rat and human NAc. **a**, Multiplexed smRNA-FISH in NAc (Blue = DAPI; scale bar = 50 µm). Magenta arrows indicate *Drd1*+/*Reln*+ cells, white arrows indicate *Drd2*+/*Reln*-cells. **b**, Left: Percent of *Drd1*+ and *Drd2*+ cells that co-express *Reln* mRNA (2-way ANOVA, interaction *p*<0.0001,). Right: *Reln* mRNA puncta by cell type (paired t-test, *p* = 0.04). **c**, Multiplexed smRNA-FISH in human NAc (Blue = DAPI; scale bar = 50 µm). Magenta arrows indicate *DRD1*+/*RELN*+ cells, white arrows indicate *DRD2*+/*RELN*-cells. **d**, Left: Percent of *DRD1*+ and *DRD2*+ cells that co-express *RELN* mRNA (paired t-test, p = 0.11). Right: *RELN* expression by cell type (paired t-test, p = 0.04)

To examine expression of *Reln* in other species, we investigated publicly available data from the Allen Institute Mouse Brain Atlas and Human Brain Atlas^49,50^. Both datasets confirmed enrichment of *Reln/RELN* mRNA expression in the adult striatum as compared to other brain nuclei (**Fig. S6**), but lacked cell type specificity in identification of *Reln/ RELN* expression. Therefore, to determine whether *RELN* is also enriched in D1-MSNs in the human brain, we performed multiplexed smRNA-FISH in post-mortem human NAc from 2 adult neurotypical donors for *RELN, DRD1*, and *DRD2*. In the human NAc, we found similar enrichment of *RELN* in *DRD1+* cells compared to *DRD2+* cells (**Fig. 2c, 2d**). Collectively, these results highlight the enrichment of mRNA for Reelin in the adult striatum, reveal that it is expressed in the principal projection neurons of the striatum, and identify conserved enrichment in D1-MSNs across species.

### CRISPR interference achieves targeted Reln knockdown in the NAc

While Reelin expression is high in the adult striatum and enriched in D1-MSNs^51^ (**Fig. 2, Fig. S5**), the molecular and cellular roles of Reelin in this region is understudied. Thus, we sought to manipulate *Reln* expression in the NAc to identify its influences on MSN physiology and drug-related behaviors. Many prior investigations of *Reln* function have employed the *Reeler* mouse model^52^, in which a large genetic deletion causes loss of *Reln* expression^45^. Similarly, mice harboring exon-skipping deletions that give rise to a truncated Reelin protein^53,54^ have been used to model similar variants observed in a schizophrenia patient^55^. However, given the germline nature of these transgenic mouse lines, the critical role Reelin plays in neurodevelopment, and the abundant expression of *Reln* outside the striatum^43,44,56^, these models do not permit specific investigation of Reelin function in the NAc. To circumvent these confounds and enable post-developmental *Reln* manipulations in a rat model system, we developed a CRISPR interference (CRISPRi) strategy to achieve targeted *Reln* knockdown in the NAc. This CRISPRi system consists of a catalytically inactive (dead) *Streptococcus pyogenes* Cas9 (dCas9) fused to the transcriptional repressor KRAB-MeCP2^57–59^ and a gRNA targeting the *Reln* promoter (**Fig. 3a-b**). As a negative control, we employed a gRNA targeting the bacterial *lacZ* gene delivered with the same dCas9-KRAB-MeCP2 (**Fig. 3a-b**). Both the *Reln* and *lacZ* gRNA vectors co-express a fluorescent mCherry reporter, enabling visualization of transduction in live or fixed cells (**Fig. 3c**). Co-delivery of these constructs using lentiviruses led to robust knockdown of *Reln* mRNA and Reelin protein in primary striatal neurons (**Fig. 3d-f**), mimicking heterozygous genetic deletions. To evaluate the specificity of this manipulation across the genome, we performed RNA-seq following *Reln* knockdown in rat primary striatal neurons (**Fig. 3g, Table S1**). As expected, we observed significant downregulation of *Reln* mRNA in neurons transduced with *Reln*-targeting CRISPRi machinery, and *Reln* was the top differentially expressed gene both by effect size (log_2_(fold change) = -2.46) and statistical significance (adjusted p value = 0). Genes overlapping with computationally identified off-target sequence predictions (allowing for sequence mismatches and RNA or DNA bulges) were not consistently or robustly altered by *Reln* CRISPRi (|log_2_(fold change)| values < 0.5; **Fig. 3g**).

**Figure 3.**
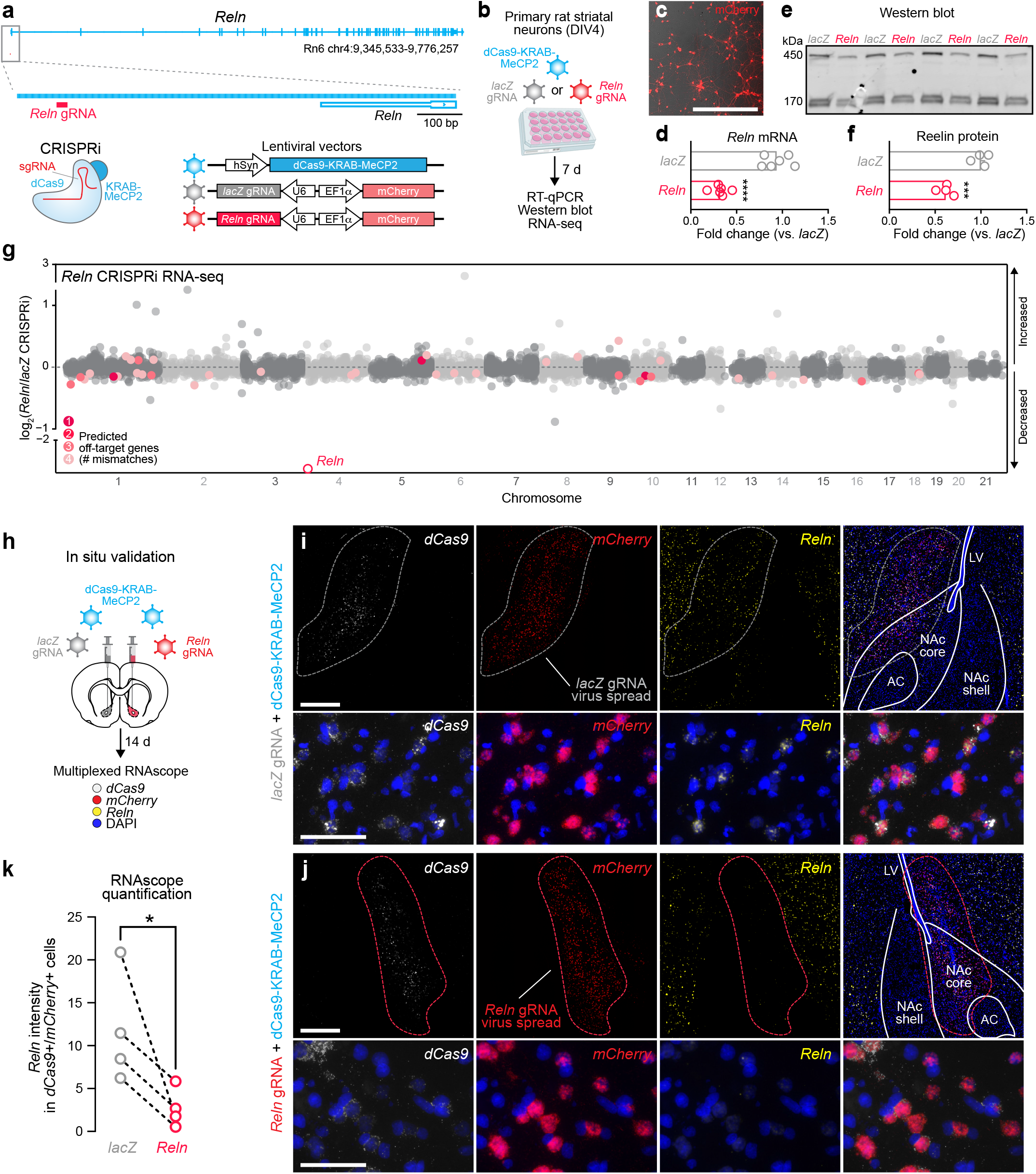
Validation of CRISPR-mediated *Reln* knockdown. **a**, Genomic location of CRISPRi *Reln* gRNA and schematic of CRISPRi lentiviral vectors for *Reln* or *lacZ* (control) targeting. **b**, Timeline for lentiviral delivery in primary striatal neurons. **c**, Represetnative image of transduction efficiency (mCherry reporter; Scale bar = 10 µm). **d**, CRISPRi at *Reln* robustly decreases *Reln* mRNA (RT-qPCR; p<0.0001, nested t-test). **e**, Western blot of secreted Reelin protein from neuronal culture media. **f**, CRISPRi at *Reln* reduces total (450 kDa + 170 kDa) Reelin protein (p <0.001, unpaired t-test). **g**, Manhattan plot highlighting signficant cas-OFFinder target predictions by number of mismatches in *Reln* CRISPRi primary striatal neuron bulk RNA-sequencing data reveals absence of off-target effects. **h**, Schematic of in vivo validation for *Reln* CRISPRi followed by smRNA-FISH. *lacZ* gRNA + dCas9 was delivered into one hemisphere and *Reln* gRNA + dCas9 was delivered into the opposite hemisphere within a single animal. **i**, Top: 40x images of the NAc (scale bar = 500 µm; dotted line indicates virus localization) of *lacZ* hemisphere. Bottom: high-magnification images (scale bar = 50 µm) showing transcript puncta from *lacZ* hemisphere. **j**, Top: 40x images of the NAc (scale bar = 500 µm; dotted line indicates virus localization) of *Reln* hemisphere. Bottom: high-magnification images (scale bar = 50 µm) showing transcript puncta from *Reln* hemisphere. **k**, In vivo CRISPRi at *Reln* successfully decreases *Reln* expression (quantification within cells expressing both *dCas9* and *mCherry* mRNA; p = 0.035, ratio paired t-test).

To assess in vivo efficacy of *Reln* CRISPRi in the NAc, we stereotaxically delivered lentiviruses expressing dCas9-KRAB-MeCP2 and *lacZ* gRNA to one NAc hemisphere and dCas9-KRAB-MeCP2 and *Reln* gRNA to the opposite NAc hemisphere (**Fig. 3h**). *Reln* knockdown was assessed using smRNA-FISH probing for *Reln, Cas9*, and *mCherry* (to identify transduced cells; **Fig. 3h-k)**. Image analysis was restricted to the region of virus spread and knockdown was quantified in cells coexpressing the gRNA and dCas9 constructs as co-transduction is necessary for knockdown. As with in vitro approaches, we detected significantly decreased *Reln* mRNA in cells targeted with CRISPRi machinery as compared to the *lacZ* control, confirming that *Reln* CRISPRi achieved robust in vivo knockdown (**Fig. 3k**).

### Loss of Reln induces transcriptional alterations in NAc D1-MSNs

To better understand Reelin’s role in the NAc, we performed snRNA-seq to identify transcriptional perturbations induced by *Reln* CRISPRi. Animals received bilateral infusions of dCas9-KRAB-MeCP2 in addition to either the *lacZ* or *Reln* gRNA. Two weeks later, NAc punches were obtained and prepped for snRNA-seq using the 10x Genomics platform as previously described^20^. Clusters were benchmarked to our published snRNAseq datasets from the NAc, with high agreement between the defined populations in this study and our previous work (**Fig. S7a**). We detected no significant differences in marker gene expression between gRNA targets in neuronal populations (**Fig. S7a**). Similarly, cell type distribution did not differ based on GEM well or treatment group (**Fig. 4c**). We verified *Reln* knockdown in D1-MSNs in the *Reln* gRNA group compared to *lacZ* controls (**Fig. 4d, Fig. S7d**), and focused subsequent analysis on this population given its selective activation by cocaine^10,20^ (**Fig. 1**). DEGs identified using rigorous pseudobulked analysis^60^ for all cell populations are noted in **Table S2**. Within the D1-MSN cluster, we identified 28 DEGs (21 upregulated and 7 downregulated; **Fig. 4e, Table S2**), with changes in several genes that function as calcium binding proteins (*Cpne4*), calcium-permeable channels (*Trpm3, Trpc5, Trpc7*), and synaptic cell adhesion proteins (*Cadm2, Cntnap4*). Gene ontology analysis for molecular function categories revealed enrichment of terms related to ion channels, led by calcium transport and signaling (**Fig. 4f**).

**Figure 4.**
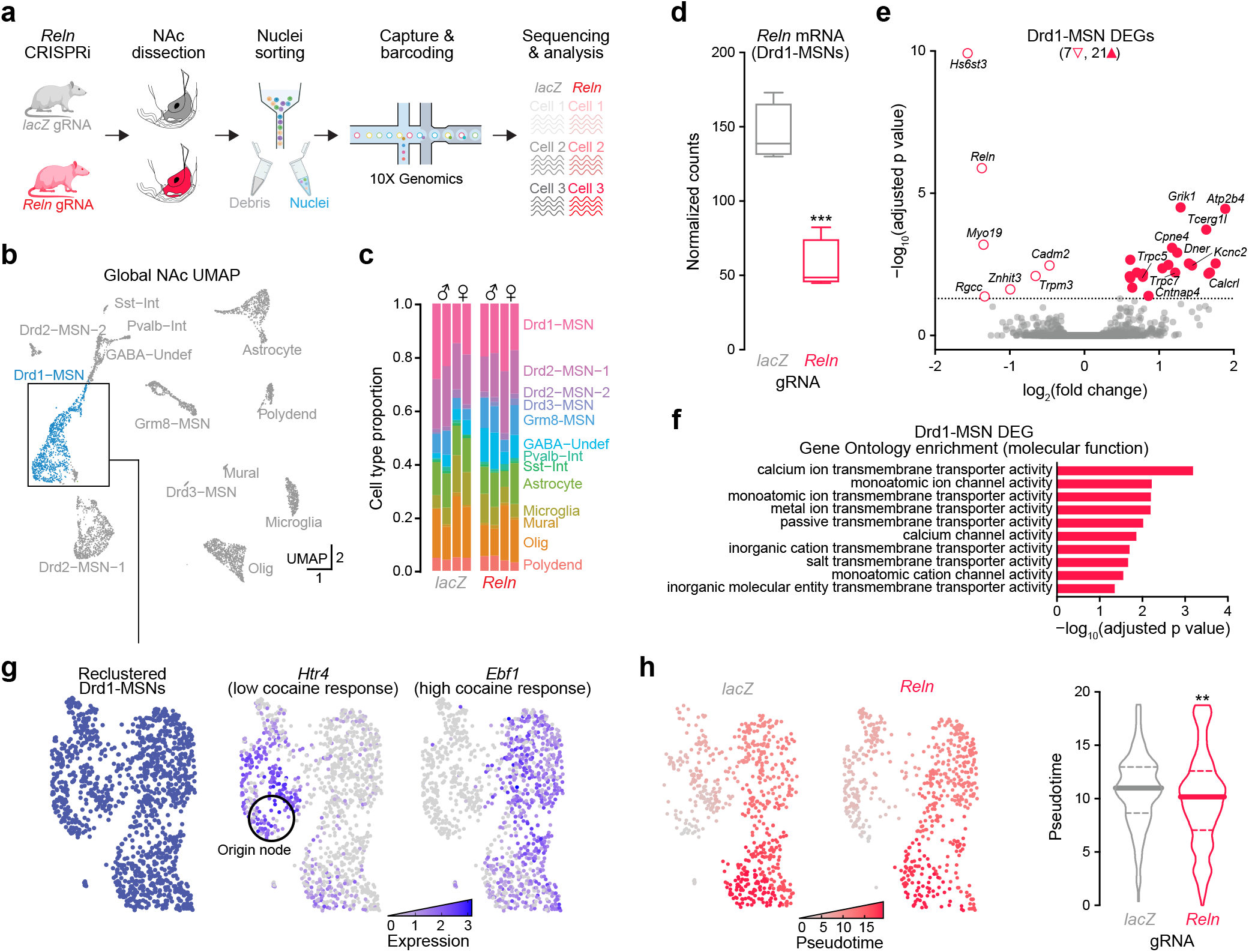
*Reln* knockdown in the NAc alters expression of calcium channel genes. **a**, snRNA-seq experimental schematic. **b**, UMAP demonstrating presence of previously described transcriptional cell types within the NAc. **c**, Cell type proportions across defined clusters is not altered by *Reln* CRISPRi. **d**, *Reln* levels are decreased in Drd1-MSNs in the *Reln* gRNA versus *lacZ* gRNA group (*p* = 0.0005, t-test). **e**, Volcano plot showing DEGs within the Drd1-MSN population. **f**, Molecular function gene ontology analysis of Drd1-MSN DEGs highlights enrichment of genes involved in calcium ion channel and transporter functions. **g**, Reclustering of Drd1-MSNs. High *Htr4* expression pocket set as origin node for pseudotime analysis. **h**, Feature plot of Drd1-MSNs colored by pseudotime score (left), with higher scores representing cell states associated with activation by cocaine. Loss of *Reln* mRNA decreased pseudotime score (right panel; Kolmogorov-Smirnov test, *p* = 0.0094).

In previous work, we demonstrated that an *Ebf1*+ D1-MSN subpopulation exhibits transcriptional sensitivity to cocaine, whereas *Htr4*+ D1-MSNs are not activated by cocaine^38^. As *Reln* marks cocaine sensitive D1-MSNs, we predicted that *Reln* knockdown would shift transcriptional profiles of D1-MSNs towards a more cocaine-insensitive state akin to *Htr4*+ D1-MSNs. To assess this possibility, we performed pseudotime analysis with the *Htr4*+ population set as the origin node (lowest pseudotime score or lightest color; **Fig. 4g, h**). To avoid the confound of *Reln* expression defining cocaine-sensitive cells, *Reln* was removed from the expression matrix and D1-MSNs were reclustered before running pseudotime (**Fig. 4g**). *Reln* knockdown led to a significant shift in the distribution of pseudotime values, with more cells having lower pseudotime values that define the putative cocaine-insensitive D1-MSN population (**Fig. 4h**). Together, these results indicate that Reelin influences MSN ion channel distribution to facilitate a cocaine-responsive state.

### Reln knockdown disrupts MSN intrinsic excitability

In addition to the induction of IEGs, cocaine also leads to persistent changes in MSN excitability^8,28^. Given that *Reln* knockdown alters the expression of various ion channels (**Fig. 4, Table S2**), we hypothesized that Reelin regulates neuronal excitability to promote a cell state necessary for cocaine responsiveness. To test this hypothesis, we performed whole-cell patch clamp electrophysiology in the NAc core following CRISPRi-mediated *Reln* knockdown (**Fig. 5a**). *Reln* knockdown had no effect on passive membrane (**Fig. 5b**) or action potential properties (**Fig. S8a-e**). To determine whether NAc-specific *Reln* knockdown impacts MSN excitability, we constructed an input-output relationship using a current step protocol. Remarkably, *Reln* knockdown led to a marked decrease in intrinsic excitability, evident by the inability of MSNs to sustain firing across increasing current steps (**Fig. 5c,d**). *Reln* knockdown cells also demonstrated a lower firing threshold during current steps (**Fig. S8f**). Additionally, MSNs exhibit a characteristic ∼200 ms first spike latency that was significantly shortened with *Reln* knockdown (**Fig. 5e**), further suggestive of altered ion channel function. Together, these results suggest that loss of *Reln* does not alter basal electrophysiological properties, but reduces MSN intrinsic excitability.

**Figure 5.**
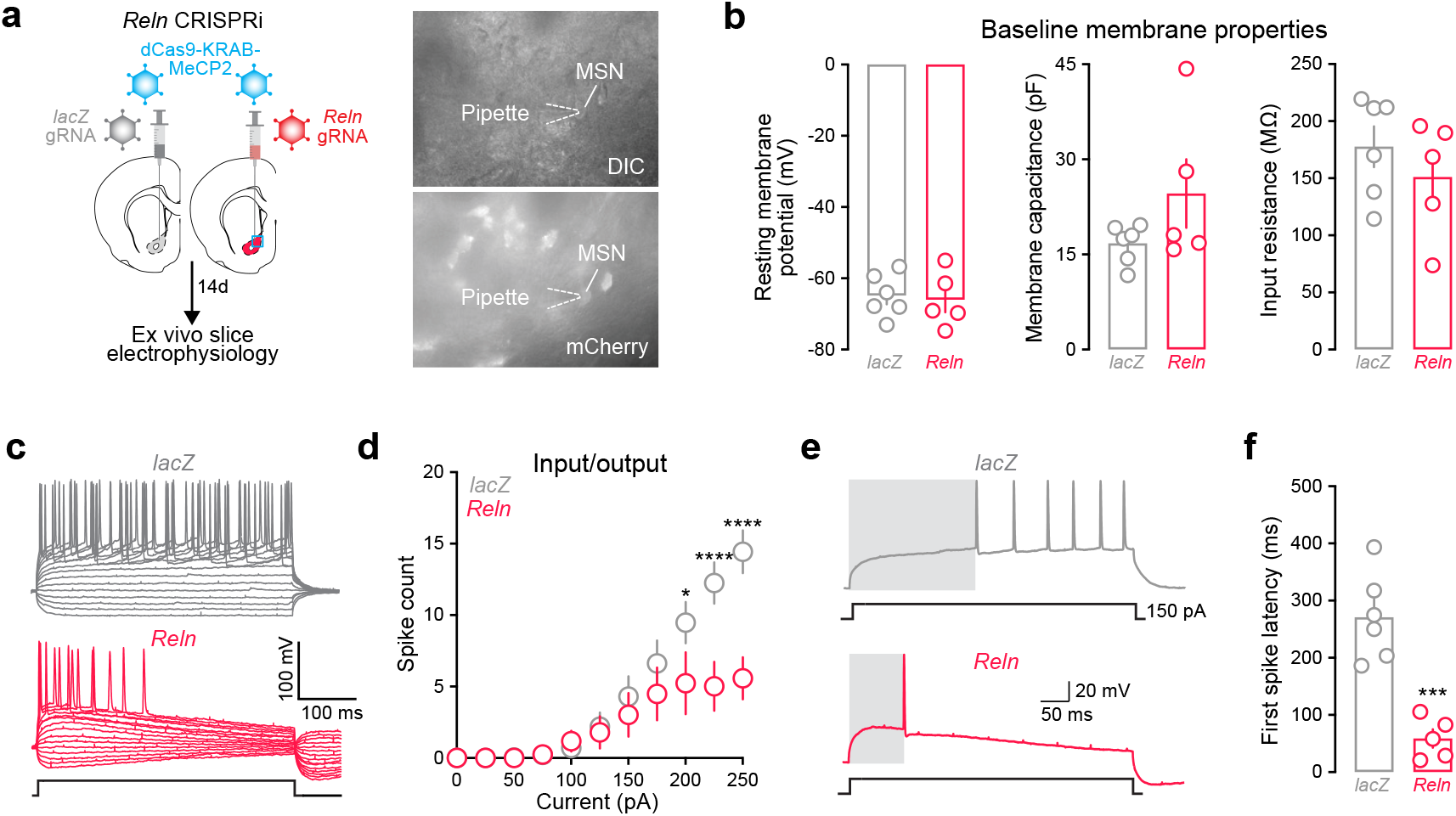
*Reln* knockdown disrupts MSN excitability. **a**, Left: Lentivirus delivery and experimental timeline. Right: Representative example of fluorescence-guided patch clamp electrophysiology in the NAc. Top: DIC image with pipette and neuron labeled. Bottom: mCherry image showing pipette recording from neuron with mCherry fluorescence. **b**, *Reln* knockdown did not alter passive properties such as resting membrane potential, membrane capacitance, or input resistance (*lacZ* n = 6 animals, 17 cells; *Reln* n = 5 animals, 11 cells; all p > 0.05, nested t-test). **c**, Representative traces across a current injection sweeps from a single representative cell from *lacZ* gRNA (gray) or *Reln* gRNA (red) recording. **d**, Input-output curve from whole-cell patch clamp electrophysiology in NAc comparing *Reln* knockdown cells to *lacZ* controls demonstrates significant attenuation of response to injected current, indicating lower intrinsic excitability (*lacZ* n = 6 animals, 17 cells; *Reln* n = 5 animals, 11 cells; p < 0.001, 2-way ANOVA with Sidak multiple comparisons). **e**, Representative traces highlighting difference in first spike latency (*lacZ* top, *Reln* bottom) at 150 pA current injection step. **f**, *Reln* gRNA neurons exhibit shortened first spike latency (*Reln* avg. latency = 59.3 ms (n = 5 animals, 11 cells) *lacZ* avg. latency = 271.2 ms (n = 6 animals, 17 cells); p = 0.0002, nested t-test).

### Reln is required for cocaine-linked behavioral adaptations

Collectively, our results suggest that *Reln* marks cocaine-activated cells and serves to promote the excitability of NAc MSNs. Given the importance of activated MSN ensembles for cocaine-related behavioral adaptations, we next sought to determine if *Reln* also contributes to the locomotor, rewarding, and reinforcing properties of cocaine. We began with a locomotor sensitization assay, a widely used paradigm in which repeated exposure to the same dose of cocaine produces an escalated locomotor response^61,62^. Two weeks following bilateral knockdown of *Reln* in the NAc (or delivery of control *lacZ* gRNA), animals received two days of i.p. saline injections, followed by two doses of cocaine (10 mg/kg, i.p.) separated by one week. We observed no differences in locomotion between *lacZ* gRNA and *Reln* gRNA animals during saline or initial cocaine exposure (**Fig. S9a,b**). While others have reported changes in anxiety-like behaviors in the setting of *Reln* deficiency^63,64^, we found no differences in either center zone entries or time spent in center during open field testing (**Fig. S9b**), indicating that *Reln* knockdown in the NAc does not lead to anxiety-like behaviors. As expected, we observed robust cocaine locomotor sensitization in the *lacZ* gRNA group, with all animals increasing locomotor response to a second dose of cocaine (**Fig. 6b**). In contrast, *Reln* gRNA animals demonstrated a lack of sensitization to the second cocaine injection (**Fig. 6b**), suggesting that loss of *Reln* prevents neuronal adaptations linked to prior cocaine history.

**Figure 6.**
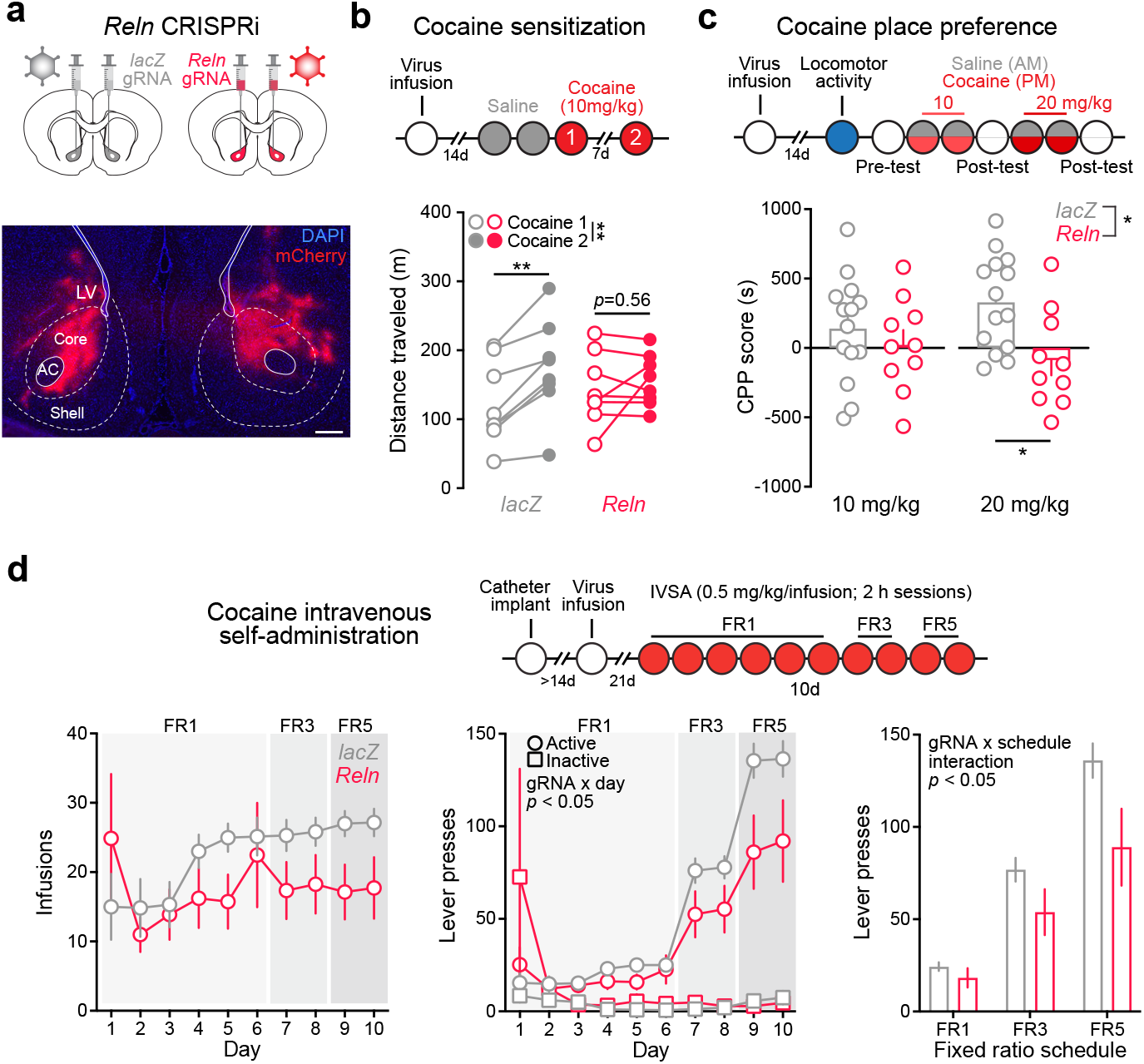
*Reln* knockdown in the NAc impairs cocaine sensitization, conditioned place preference, and self-administration without altering locomotion. **a**, Top: schematic of bilateral lentiviral targeting; bottom: representative image showing expression of mCherry reporter (from gRNA lentivirus) in the NAc (scale bar 500 µm). **b**, *Reln* knockdown in the NAc prevents locomotor sensitization to cocaine (2-way repeated measures ANOVA; p = 0.0465 for gRNA x time interaction). **c**, *Reln* knockdown abolishes conditioned place preference for cocaine (CPP score = time spent in cocaine paired chamber - time spent in saline paired chamber; 2-way ANOVA with mutliple comparisons and Tukey’s correction; p = 0.034 for main effect of gRNA, p = 0.014 for effect of gRNA at 20 mg/kg). **d**, Effects of *Reln* knockdown on cocaine IVSA infusions (left panel) and lever presses (center and right panels). Across days, *Reln* knockdown led to significantly fewer active lever presses (2-way ANOVA for active presses, p = 0.0009 for interaction between gRNA and days). *Reln* knockdown resulted in a schedule-dependent decrease in cocaine lever presses (2-way ANOVA, p = 0.0472 for interaction between gRNA and FR schedule).

While locomotor sensitization is frequently used as a behavioral correlate of cocaine-induced molecular and cellular plasticity, it does not directly measure cocaine reward or memory. To assess this facet of behavior, we used the cocaine conditioned place preference paradigm (CPP; **Fig. 6a,c**; **Fig. S9c,d**). Two weeks following viral infusion of *Reln* CRISPRi vectors into the NAc, we performed an open field assay and found no differences in baseline locomotion nor anxiety-like behaviors, as in our locomotor sensitization cohorts (**Fig. S9c,d**). Next, we performed cocaine CPP pairings at 10 and 20 mg/kg doses, each followed by a post-test session which enabled quantification of place preference. *Reln* CRISPRi produced lower preference for the cocaine-paired compartment, with an overall main effect of *Reln* gRNA and complete reversal of cocaine CPP at 20 mg/kg (**Fig. 6d**). These results demonstrate that selective knockdown of *Reln* in the NAc alters reward-related behavior without impacting locomotion or anxiety-like behavior.

We next investigated if effects of *Reln* knockdown extend to intravenous self-administration (IVSA), asdrug context (i.e., experimenter administered versus self-administered) can affect the cellular and behavioral response to psychostimulants^65–67^. Jugular vein catheterized animals received bilateral infusions of lentiviruses expressing dCas9-KRAB-MeCP2 and either *lacZ* or *Reln* gRNA three weeks prior to beginning cocaine IVSA acquisition and responding on a fixed ratio schedule (**Fig. 6a,d; Fig. S9e**). Both groups exhibited stable self-administration behavior, with expected sensitivity to changes in reinforcement schedule requirements (**Fig. 6d**). Control (*lacZ*) rats escalated cocaine infusions across behavioral training sessions (significant linear correlation between acquisition day and infusion number; R^2^ = 0.295, *p* < 0.0001), whereas this behavior was absent in the *Reln* CRISPRi group (R^2^ = 0.0001, *p* = 0.907). Additionally, loss of *Reln* decreased lever pressing behavior in a schedule-dependent way, with larger effects observed when increased effort was required for each cocaine infusion (**Fig. 6d**). Together, these results suggest that CRISPR-mediated *Reln* knockdown decreased the sensitizing, rewarding, and reinforcing properties of cocaine, without altering overall locomotion or initial locomotor response to cocaine.

## DISCUSSION

Drug-activated ensembles are key mediators of the cellular and behavioral effects of drug experience^10^. The molecular changes that occur downstream of striatal neuron activation are increasingly well understood, and include activation of signal transduction cascades, chromatin and transcriptional reprogramming, and coordinated alterations in expression of key receptors, neuropeptides, and ion channels^17,20,68–72^. Despite these advances, determinants of ensemble participation have previously remained elusive. In this study, an unbiased approach identified *Reln* as a marker of a cocaine-activated neuronal ensemble in the NAc. While *Reln* is not altered after cocaine experience, it is abundantly expressed in NAc neurons expressing IEGs following cocaine administration, suggesting that *Reln* plays an active role in MSN excitability. To test this prediction, we developed a CRISPRi approach to repress *Reln* expression from the endogenous gene locus. NAc-specific loss of *Reln* disrupted transcriptional patterns in D1-MSNs, impaired intrinsic excitability of MSNs, and decreased cocaine-related behavioral adaptations. These results establish *Reln* as both a marker and a regulator of cocaine-induced plasticity in the NAc.

In the developing brain, Reelin plays a central role in cortical layering^44,45^ and segregation of the dopaminergic system^73,74^. Postnatally, Reelin regulates synaptic function and long-term potentiation in the hippocampus through both post-translational^43,75,76^ and transcriptional mechanisms^46^. Additionally, Reelin has links to diverse neuropsychiatric disorders, such as schizophrenia^53,77–79^, substance use disorders^80,81^, and Alzheimer’s disease^82,83^. Secreted Reelin protein is cleaved in the extracellular matrix by a variety of metalloproteases^84^, and canonically acts at low-density lipoprotein receptors such as LRP8 (also known as ApoER2) and VLDLR^43^, although non-canonical modes of action have also been identified^85^. Reelin binding at low-density lipoprotein receptors initiates cell signaling cascades largely mediated by the intracellular adapter protein DAB1, including phosphorylation of NMDA receptors by Src and Fyn family kinases and activation of the PI3K-Akt-mTor pathway^43,56^. Despite the presence of all of this machinery in the adult striatum and close links between Reelin and diseases arising from striatal dysfunction^53,77,78,80^, few studies have explored Reelin’s role in basic striatal biology. This lack of investigation likely stems from difficulties associated with measuring adult phenotypes in a model lacking a critical neurodevelopmental protein. Our novel CRISPRi approach avoids these developmental confounds, and also enabled site-specific reduction of *Reln* in the NAc.

Well-established models of Reelin function hold that after the disappearance of Reelin-expressing Cajal-Retzius cells during neurodevelopment, Reelin levels are principally maintained by GABAergic interneurons^43^. Therefore, the discovery that *Reln* mRNA marks projection neurons (MSNs) in the NAc that respond to cocaine was surprising. However, this finding is in line with recent studies showing high expression of *Reln* in the murine dorsal and ventral striatum, as well as localization within MSNs^51,86,87^. In agreement with our findings, these studies also identified *Reln* enrichment in *Drd1*+ cells as compared to *Drd2*+ cells in the mouse dorsomedial striatum^51^, and higher expression in the NAc core versus shell^87^. Our results build on these findings to characterize *Reln* expression throughout the entire striatum, confirming enrichment in *Drd1*+ versus *Drd2*+ populations across striatal subregions in the rat brain (**Fig. S5**). We further define a dorsoventral and mediolateral gradient of *Reln* expression in the striatum, with the highest *Reln* levels in the dorsolateral striatum and near absence of *Reln* in the ventromedial NAc shell (**Fig. S5**). Additionally, we validate the abundant expression of *RELN* mRNA in the human NAc, and also reveal enrichment in *DRD1*+ human neurons. These findings align closely with large scale transcriptional profiling studies of the human NAc^88^, and highlight the translational relevance of *Reln* perturbations in animal models.

Our results define a previously undescribed role for Reelin in MSN physiology, with loss of *Reln* leading to a striking decrease in intrinsic excitability and altered expression of ion channels and calcium signaling proteins. This observation builds upon a strong body of literature supporting plasticity-promoting functions of Reelin in the hippocampus^75,76,89^. Recent work in hippocampal neurons has also linked Reelin to neuronal excitability, via NMDA-dependent regulation of calcium-permeable AMPA receptors^90^. Similarly, synaptic Reelin signaling gates ketamine-induced plasticity in the hippocampus via maintenance of basal neurotransmission at the NMDA receptor^91^. However, in the hippocampus *Reln* mRNA is expressed by local interneurons, and is absent in pyramidal neurons^92,93^. Therefore, it is likely that the relationship between Reelin secretion and excitability in the striatum may be distinct from prior observations in the hippocampus, and future work will be required to comprehensively delineate these differences.

Rapid induction of IEGs such as *Fos* and *Arc* is a hallmark of electrical or synaptic activation across the nervous system^94–96^. This relationship has been leveraged to generate a variety of widely used tools enabling permanent tagging and functional manipulations within neuronal ensembles^97–100^. While the critical roles of various IEG transcription factors in plasticity of different brain circuits are increasingly appreciated^69,101–103^, considerably less is known about the mechanisms that regulate ensemble participation. Foundational work established that regulation of neuronal excitability by the activity-responsive transcription factor CREB served as a marker of activated neuronal ensembles^40–42,104,105^. However, these studies all focused on activated ensembles in the amygdala, and similar markers of activated striatal ensembles have not been identified or functionally tested. The present results provide robust evidence from orthogonal and unbiased approaches that cocaine results in IEG transcription in a small population of *Reln*+ neurons in the NAc. While our findings provide a mechanistic link between Reelin expression and ensemble recruitment, it is unclear if Reelin similarly marks activated ensembles in other brain structures (or in response to other drugs of abuse or other rewarding stimuli). Drugs of abuse and natural rewards engage distinct yet partially overlapping NAc ensembles^18,106^. Ensemble participation is partially determined by drug class, with cocaine (a stimulant) primarily engaging D1-MSNs^10,20^ and morphine (an opioid) recruiting both D1- and D2-MSNs^10,107^. However, cocaine does activate a small group of D2-MSNs^9,20^, and we similarly find enrichment of *Reln* mRNA in this MSN population (**Fig. S3**). Thus, it is likely that *Reln* marks ensembles that span the boundaries between broad cell classes, and may also mark cells that respond to other reward or drug experiences.

In addition to identifying *Reln* as a marker of a cocaine-activated ensemble, we report that loss of NAc *Reln* abolishes cocaine sensitization and place preference memory, and decreases cocaine self-administration behavior. Prior studies investigating the role of Reelin in psychostimulant responses have generated mixed and often conflicting results. For example, haploinsufficient *Reeler* mice or mice harboring loss-of-function Reelin truncations exhibit decreased locomotor sensitivity to methamphetamine^53,108^. However, *Reeler* mice display a normal locomotor response to amphetamine^109^, and increased locomotor activity in response to cocaine, with intact cocaine CPP^51^. Likewise, transgenic forebrain overexpression of *Reln* using the *Camk2a* promoter decreased cocaine locomotor sensitization without altering initial cocaine response^77^. Given the strong developmental role of *Reln*, as well as its important functions in brain regions that send crucial projections to the NAc, these divergent results emphasize the need for continued exploration of Reelin’s role in the adult striatum.

Reelin’s necessity in cocaine-related behavioral adaptations supports the notion that cocaine ensembles consist of a primed population of neurons whose activation is crucial for diverse cocaine-related adaptations. Consistent with this idea, inactivation of cocaine-sensitive ensembles (identified based on IEG expression) ablates or dampens subsequent cocaine-related behaviors, defining a key role for ensembles in adaptations to future exposure. Here, we identify *Reln* as a stable marker of this ensemble, which may facilitate future efforts for *a priori* tagging, monitoring, and manipulation of this population. Remarkably, our work also suggests that *Reln* does not simply serve as a marker of this ensemble, but likely plays an essential role in promoting a cocaine-sensitive state. Taken together, these findings highlight an opportunity for high-precision manipulation of reward circuitry using *Reln*-based tools, reveal the necessity of Reelin in the cellular response to cocaine, and implicate the Reelin signaling pathway as a potential therapeutic target for cocaine use disorder.

## METHODS

### Animals

All experiments were performed in accordance with the University of Alabama at Birmingham Institutional Animal Care and Use Committee. Sprague-Dawley adult male and female rats (45 days old, patch clamp experiments; 90-120 days old / 225-250g all other experiments) were purchased from Charles River Laboratories (Wilmington, MA, USA). Rats were co-housed in pairs in plastic filtered cages with wooden chewing block enrichment in an AAALAC-approved animal care facility maintained between 22-24°C on a 12-hour light/dark cycle with ad libitum access to food (Lab Diet SL3Z Irradiated rat chow) and water. Bedding and enrichment were changed weekly by animal resources program staff (no changes during behavioral testing as to not disturb animals). Animals were randomly assigned to experimental groups. All animals were handled by investigators for 5-7 days prior to behavioral testing.

### Drugs

Cocaine hydrochloride (C5776, Sigma-Aldrich, St. Louis, MO, USA) was dissolved in sterile 0.9% sodium chloride and injected intraperitoneally (i.p.) at described doses (10 or 20 mg/kg) for cocaine conditioned place preference testing, cocaine locomotor response, and smRNA-FISH studies. For smFISH studies, cocaine was prepared as in behavioral testing at a dose of 20 mg/kg for i.p. injections. For IVSA, cocaine was dissolved in sterile 0.9% sodium chloride for a final concentration of 2.5 mg/kg/mL (or 0.5 mg/mL/infusion). Two separate solutions were prepared for each sex based on the average weight of each sex. Cocaine solution was made fresh before behavioral testing and was protected from light.

### Reln gRNA construction

The *Reln* gRNA was designed using an online sgRNA tool, provided by the Zhang Lab at MIT (crispr.mit.edu) and inserted in a previously described lentivirus compatible sgRNA scaffold construct. To ensure specificity, all CRISPR crRNA sequences were analyzed with the National Center for Biotechnology Information’s (NCBI) Basic Local Alignment Search Tool (BLAST) and Cas-OFFinder (http://www.rgenome.net/cas-offinder/).

### Lentivirus production

Lentiviruses were produced in accordance with BSL-2 safety guidelines in a biosafety cabinet, as previously described. ^20^ For in vivo preparations, HEK293T cells were triple transfected with lentivirus packaging plasmid psPAX2 and the envelope plasmid pCMV-VSV-G (Addgene plasmids #12260 and #8454) in addition to the transgene of interest using FuGene HD (Promega). Media was harvested at 48- and 72-hours post-transfection. Following harvest, media was filtered (0.45 µm) and virus pelleted via ultracentrifugation (25,000 RPM, 1 hr 45 min) and then resuspended in phosphate--buffered saline. Titers were obtained on freeze-thawed aliquots using the qPCR-based TaKaRa lentivirus titration kit (#631235, Takara Bio, Kusatsu, Shiga, Japan). Minimum titers for all in vivo experiments was 1×10^11^ genome copies/mL. Virus was stored in single-use aliquots at -80°C. Prior to surgical infusion, viruses were thawed on wet ice and combined such that an equivalent number of genome copies for gRNA and effector made up the final mixture.

### Stereotaxic Surgery

Naïve adult male and female Sprague-Dawley rats (Charles River) were anesthetized with 4-5% isoflurane and placed in a stereotaxic apparatus (Kopf instruments, Tujunca, CA, USA). Rats were maintained at a surgical plane of anesthesia with 2-3% isoflurane, and respiratory rates were monitored throughout surgery and maintained between 35-55 respirations per minute. Surgical coordinates were determined using the Paxinos and Watson rat brain atlas (6^th^ edition), targeting the NAc core bilaterally in *Reln* knockdown validation and behavioral studies and unilaterally for patch clamp experiments. Under aseptic conditions, guide holes were drilled at AP + 1.7mm, ML ± 1.5mm, and the infusion needle was lowered to DV – 7.2mm (all coordinates with respect to bregma^110^). All infusions were made using a gastight 30-gauge stainless steel injection needle and 10 µL syringe (Hamilton Company, Reno, NV, USA). Lentivirus constructs were infused bilaterally at a rate of 0.25 µL/min using a syringe pump (Harvard Apparatus, Holliston, MA, USA), totaling 1.5 µL/hemisphere. Following each infusion, needles remained in place for 10 minutes to allow for diffusion of the virus. Infusion needles were slowly retracted and guide holes were filled with sterile bone wax. The surgical incision was closed with interrupted 6-0 monofilament nylon sutures (Covidien, Dublin, Ireland). At the end of surgery, rats were administered buprenorphine (0.03 mg/kg) and carprofen (5 mg/kg) for analgesia, and topical bacitracin (500 units) was applied to the incision site as a skin protectant and antimicrobial treatment. Animals were placed on a heating pad covered by a sterile drape to maintain body temperatures during surgery.

### Acute cocaine injections

For in situ validation of *Reln* as a marker for cocaine sensitive D1 neurons, animals (n = 2/sex/group) received a homecage i.p. injection of either 20 mg/kg cocaine or saline. Animals were euthanized 1 h following injection, and tissue was harvested for smRNA-FISH as described below.

### Locomotor sensitization

Two weeks following viral infusion surgeries, animals (n = 4 animals/sex/ group) received i.p. saline injections on Days 1 and 2 immediately prior to locomotor testing in an open field chamber for 30 minutes. On Day 3, animals received 10 mg/kg cocaine i.p. (equivalent in volume to saline injections) prior to locomotor testing. One week later (Day 10), animals received another 10 mg/kg i.p. cocaine injection followed by 30 minutes of locomotor testing. All locomotor activity was monitored in a 43 cm × 43 cm plexiglass locomotor activity chamber (Med Associates, Inc., St. Albans, VT, USA) with opaque white wall covering and an open top.

### Baseline locomotor testing

Prior to conditioned place preference testing, baseline locomotor (n = 4 animals/ sex/group) activity was monitored for 30 minutes as described for locomotor sensitization experiments.

### Conditioned place preference testing

Conditioned place preference (CPP) testing was completed in a three-chamber apparatus with guillotine-style doors (Med Associates, Inc.) using previously established protocols with minor modifications ^72^. The two chambers used for conditioning measured approximately 27 x 21 x 22 cm, one with opaque black walls and stainless steel bar flooring, and the other with opaque white walls and metal stainless steel grid flooring. The conditioning chambers were separated from one another by manual guillotine-style doors and a central gray compartment with solid flooring, measuring 12 x 21 x 22 cm. All three chambers had clear perforated acrylic lids with centrally-placed house lights. Each apparatus included a 16-channel infrared controller (Med Associates, Inc.) to track rodent position, in conjunction with Med PC software (v4.1.49; Med Associates, Inc.). CPP testing began 2 weeks following lentiviral infusion surgeries of CRISPRi machinery targeting the NAc core. Male and female rats (n =14 *lacZ* (6 M/8 F) *Reln* (6 M/4 F) were placed in the central compartment on the first day of testing (i.e., pre-test) and were permitted to explore all three chambers of the CPP apparatus during a 30-minute session. Days 2 and 3 of testing were conditioning days. In the morning sessions, rats were given an i.p. injection of saline immediately before being placed in the initially preferred chamber for the 30-minute session. In the afternoon sessions, rats were given an i.p. injection of 10 mg/kg cocaine prior to placement in the other (initially non-preferred) chamber. A post-test for the 10 mg/kg dose was performed on Day 4, where animals had free access to all 3 chambers as in the pre-test on Day 1. Animals then went through 2 more pairing days (Days 5 & 6) for 20 mg/kg cocaine with the post-test on Day 7.

### Intravenous self-administration

Intravenous self-administration was completed in an operant conditioning box (Med Associates Inc), measuring 29.5 x 23.5 x 27.3 cm with retractable response levers, a house light and fan, cue lights above each lever, and a syringe pump located outside of a sound-attenuating chamber that surrounded the operant box. Jugular vein catheterization surgeries were performed by the vendor (Charles River) prior to arrival at the University of Alabama at Birmingham, and 22 gauge vascular access buttons (Instech, VABR1B/22) were used for intravenous access. Cocaine IVSA began 3 weeks following bilateral intracranial infusions for CRISPRi lentiviruses targeting the NAc core (n = 6 *lacZ* (2 M/4 F), 8 *Reln* (4 M/4 F). Sessions started using a computer--issued command, signaled by illumination of the house light, fan, and protraction of both inactive and active levers. Responses on the active lever resulted in illumination of a cue light above the lever and lever retraction for 20 s, as well as drug infusion (infusion time 2.5 s, infusion rate 2.636 mL/min). Responses on the inactive lever had no programmed consequences. All animals began on a fixed ratio (FR) 1 schedule reinforcement for each cocaine infusion (0.5 mg/kg/ infusion), which was maintained for the first 6 days of IVSA. FR schedules were then increased to FR3 (for 2 days), and then FR5 (for 2 days). All sessions lasted for 2 h or until 100 infusions were reached.

### Tissue collection from adult rat Nac

Animals were rapidly decapitated using a guillotine (World Precision Instruments, Sarasota, FL, USA). Brain tissue was extracted and submerged in cold 2-methylbutane, chilled on dry ice, for approximately 1 minute. Flash-frozen brains were then removed from the 2-methylbutane, wrapped in aluminum foil, and placed on dry ice. Tissue was stored at -80°C and equilibrated to -20°C for at least one hour prior to sectioning at -18°C on a Leica CM 1850 cryostat (Deer Park, IL, USA) in 10µm sections.

### smRNA-FISH and image acquisition in fresh-frozen rat tissue

smFISH was performed (#323136, ACD Bio, Newark, CA, USA) to probe for *Reln, Drd1* and *Fos* (*Fos* quantification) or *Reln, Drd1*, and *Drd2* (*Reln* characterization) or *Reln, mCherry*, and *dCas9* (*Reln* knockdown) following manufacturer’s protocol for fresh-frozen tissue, as previously described^111 112^. Briefly, tissue was submerged in ice-cold 10% normal-buffered formalin for 15 minutes and then serially dehydrated with ethanol, then treated with hydrogen peroxide, followed by protease IV digestion. Slides were incubated with a combination of 3 probes (Characterization study: *Reln, Drd1, Drd2*; *Fos* quantification: *Reln, Drd1, Fos*; *Reln* knockdown: *Reln, mCherry, dCas9* (#1048921-C4, #317031-C3, #315641, #403591-C2, #43120, #519411-C2; ACD Bio, Newark, CA, USA) and stored overnight in a 5x saline-sodium citrate (SSC) buffer. Probes were fluorescently labeled with Opal Dyes (Akoya Biosciences, Marlborough, MA, USA; Characterization: Opal 520 assigned to *Drd1*, Opal 570 assigned to *Reln*, Opal 690 assigned to *Drd2*; *Fos* Quantification: Opal 520 assigned to *Drd1*, Opal 570 assigned to *Fos*, Opal 690 assigned to *Reln*; *Reln* knockdown: Opal 520 assigned to *spCas9*, Opal 570 assigned to *mCherry*, Opal 690 assigned to *Reln*; all dyes diluted 1:750). Sections were stained with DAPI (4’,6-diamidino-2-phenylindole) and cover-slipped with ProLong Glass Antifade Mountant (P36984, Thermo Scientific, Waltham MA, USA). All images were acquired on a Keyence-BZ800 microscope. Within an experiment, all images were acquired with the same acquisition settings for a given objective and channel. For *Reln* characterization studies, 20x images of one striatal hemisphere were obtained across 8 animals. Images were stitched using Keyence Image Analyzer software. For *Reln* knockdown studies, 20x images were obtained of the whole striatum and stitched using the Keyence Image Analyzer Software. For *Fos* quantification studies, 40x images of one striatal hemisphere were obtained per animal and stitched using the Keyence Image Analyzer software. For regional analyses of stitched images, rat brain atlas overlays were applied to the stitched images and used to make regional crops in Adobe Photoshop (Paxinos & Watson, 6^th^ edition). For puncta analysis in *Reln* characterization studies, 8-14 images per striatal subregion per animal were obtained. For *Fos* quantification studies, 100x images of the NAc were obtained and stitched using Keyence Image Analyzer. All 100x images were converted to Nikon (.nd2) file formats and puncta counts were obtained using QuPath software (version 0.3.2).

### Rat smRNA-FISH image analysis

Images were thresholded and masked in ImageJ. Thresholds were obtained by taking the average value of thresholds for each image in a given channel. DAPI regions of interest (ROIs) were created using the StarDist-2D plugin (“Versatile fluorescent nuclei”) mode, probability score of 0.45, overlap threshold of 0.20). These DAPI ROIs were then applied to the different channels corresponding to *Reln, Drd1, Drd2*, or *Fos* which were converted to 8-bit images and thresholded based on an averaged auto-threshold value specific for each channel. Positivity was identified based on mean intensity for each channel in the DAPI ROIs. Data was further processed using R to identify signal overlap for individual DAPI ROIs to identify cell types. For *Reln* knockdown quantification, *Reln* intensity scores were used to quantify *Reln* expression in cells that co-expressed *mCherry* (gRNA) and *dCas9*. For puncta analysis, thresholds were defined in QuPath v0.3.2 ^113^ and held consistent for each channel across regions and animals. In downstream analyses, K-means clustering was used to identify relevant threshold values for puncta counts. R was again used to identify cell types based on the overlap of puncta within a given nuclear ROI.

### Post-mortem human tissue samples

Postmortem human brain tissue from two neurotypical adult donors was obtained at the time of autopsy through the Office of the Chief Medical Examiner of the State of Maryland following informed consent from legal next-of-kin, under the Maryland Department of Health IRB protocol #12-24. Clinical characterization, diagnoses, and macro- and microscopic neuropathological examinations were performed on all samples using a standardized paradigm, and subjects with evidence of macro- or microscopic neuropathology were excluded. Details of tissue acquisition, handling, processing, dissection, clinical characterization, diagnoses, neuropathological examinations, RNA extraction and quality control measures have been described previously^114^. The nucleus accumbens was dissected using a hand held dental drill at a level where the caudate and putamen are joined by the accumbens at the ventral aspect of the striatum, with clear striations separating the putamen from the caudate. The anterior aspect of the temporal lobe and the claustrum served as additional landmarks.

### smRNA-FISH and image acquisition in post-mortem human tissue

Two blocks of fresh frozen NAc from neurotypical control individuals were sectioned at 10μm and stored at −80°C. For post-mortem human studies (**Figure 2c,d**), in situ hybridization assays were performed with RNAscope technology utilizing the RNAscope Fluorescent Multiplex Kit V2 and 4-plex Ancillary Kit (Cat # 323100, 323120 ACD, Hayward, CA, USA) according to manufacturer’s instructions. Briefly, tissue sections were fixed with a 10% neutral buffered formalin solution (Cat # HT501128 Sigma-Aldrich, St. Louis, MO, USA) for 30 min at RT, series dehydrated in ethanol, pretreated with hydrogen peroxide for 10 min at RT, and treated with protease IV for 30 min. Sections were incubated with probes for *RELN, DRD1*, and *DRD2* (Cat# 413051-C2, 553991-C3, 524991-C4, ACD, Hayward, CA, USA) and stored overnight in 4x SSC buffer. Probes were fluorescently labeled with Opal Dyes (Perkin Elmer, Waltham, MA; Opal570 diluted at 1:1500 and assigned to RELN; Opal690 diluted at 1:500 and assigned to DRD2; Opal620 diluted at 1:500 and assigned to *DRD1*) and stained with DAPI to label the nucleus. Images were acquired in z-series using a Zeiss LSM780 confocal microscope as previously described ^115^.

### Human smRNA-FISH image analysis

Automated image processing was performed using the *dotdotdot* MATLAB-based command line toolbox^115^, which is available at https://github.com/LieberInstitute/dotdotdot. Downstream analyses were performed using R v4.0.0+2020.04.24.

### Neuronal cell cultures

Primary rat striatal cell cultures were generated from E18 striatal tissue as described previously^20,57,71^. Cell culture plates (Denville Scientific, Inc., Plainfield, NJ, USA) were coated overnight with poly-L-lysine (50 µg/mL; Sigma-Aldrich), supplemented with 7.5 µg/mL laminin (Sigma-Aldrich), and rinsed with diH_2_O. Dissected striatal tissue was incubated with papain (LK003178, Worthington Biochemical Corporation, Lakewood, NJ, USA for 25 min at 37°C. After rinsing in complete Neurobasal media (supplemented with B27 and L-glutamine, Invitrogen), a single-cell suspension was prepared by sequential trituration through large to small fire-polished Pasteur pipettes and filtered through a 100 µm cell strainer (Fisher Scientific, Waltham, MA, USA). Cells were pelleted, re-suspended in fresh media, counted, and seeded to a density of 125 000 cells per well on 24-well culture plates (65 000 cells/cm^2^). Cells were grown in complete Neurobasal media for 12 days in vitro (DIV12) in a humidified CO_2_ (5%) incubator at 37°C with half-media changes at DIV1, 4-5, and 8-9.

### RNA extraction and qPCR

RNA was extracted per manufacturer’s instructions using the Qiagen RNeasy RNA extraction kit (#74106, Qiagen, Hilden, Germany). cDNA libraries were synthesized using iScript Supermix (Bio-Rad). Expression of *Gapdh* (Forward: ACCTTTCATGCTGGGGCTGGC; Reverse:GGGCTGAGTTGGGATGGGGAC) and *Reln* (Forward: AATCGGCAAGAGTCACCGAAGCC; Reverse: CTGTCTGTGCCGTGCTCCCAA) were measured using RTqPCR with the SYBR Green detection.

### Bulk RNA-sequencing

Primary rat striatal neurons were transduced with lentiviruses expressing CRISPRi machinery on DIV12 with a multiplicity of infection (MOI) per cell of 2000 for each virus. Primary neurons were harvested one week later on DIV19 after which RNA was extracted (RNeasy, QIAGEN) and submitted to the Genomics core lab at the Heflin Center for Genomic Sciences at the University of Alabama at Birmingham for library preparation. Four biological replicates were used per gRNA. 100 bp paired-end libraries were sequenced on an Illumina NovaSeq 6000 instrument. Raw FASTQ files were processed using nf-core/rnaseq v3.10 (doi: https://doi.org/10.5281/zenodo.1400710) of the nf-core collection of workflows^116^ using a pipeline executed with Nextflow v23.04.1^117^. Splice-aware alignment to the mRatBn7.2/Rn7 genome assembly was conducted using STAR v2.7.9a ^118^ with the associated Ensembl gene transfer format (gtf) file (version 105). Binary alignment map files were indexed with SAMtools^119^ (v1.16.1). Gene-level counts were generated using featureCounts (Rsubread^120^ v2.0.1). QC metrics were collected and reviewed with MultiQC^121^ (v1.13), and a minimum of 20M mapped reads were obtained per sample (range: 22.6M to 121.4M). Differential expression testing was conducted using DESeq2^122^ (v1.38.3). DEG testing p-values were adjusted with the Benjamini–Hochberg method ^123^, and DEGs were designated as those genes with an adjusted p-value <0.05. Genes with unreliable/low expression estimates (basemean <50) were excluded from subsequent analysis.

### Off-target detection for CRISPRi

Potential off-target sites for CRISPRi were identified using Cas-OFFinder^124^, allowing up to 2 DNA/ or RNA bulges and up to 4 mismatches. Overlap between DEGs and CasOFFinder hits were assessed using the IRanges and GenomicAlignments tools with findOverlap() function^125^. Manhattan plot highlighting lack of off-target effects was generated in RStudio using the Manhattan package available at https://github.com/boxiangliu/manhattan/.

### Western blot

Media was harvested from rat primary striatal neuron cultures on DIV19, seven days following transduction with lentiviruses expressing CRISPRi machinery to target *lacZ* or *Reln*. 50 µL of media was reduced and denatured prior to gel electrophoresis on a 4-15% tris-glycine gel (#4561084, Bio-Rad Laboratories, Hercules, CA, USA) for 2 hours @ 175 mV. Protein was transferred to a PVDF membrane at 4°C for 16 hours at 100 mV. Membranes were incubated in monoclonal Reelin antibody (Millipore Cat# MAB5364, <u>RRID:AB_2179313</u>) overnight at 4 degrees before incubation with secondary for 2 hours (#926-68072, Li-Cor Biosciences, Lincoln, NE, USA) at room temp. Blots were imaged on a Li-Cor Odyssey 9120. Protein was quantified in ImageJ using the gel analyzer tool to obtain area under the curve.

### Whole-cell patch clamp electrophysiology

To examine the role of Reelin in intrinsic excitability, whole-cell patch-clamp recordings were conducted as previously described ^126^. Rats 60-120 days old were briefly anesthetized with isoflurane and intracardially perfused with oxygenated ice cold recovery solution: (in mM) 93 NMDG, 2.5 KCl, 1.2 NaH_2_PO_4_, 30 NaHCO_3_, 20 HEPES, 25 glucose, 4 sodium ascorbate, 2 thiourea, 3 sodium pyruvate, 10 MgSO_4_(H_2_O)_7_, 0.5 CaCl_2_(H_2_O_2_), and HCl added until pH was 7.3-7.4 with an osmolarity of 300-310 mOsm. Coronal slices (300 µm) were prepared on a vibratome (Leica VT1200S) containing recovery solution and then transferred to a Brain Slice Keeper (Automate Scientific) containing a holding solution for at least 1 hour prior to recording: (in mM) 92 NaCl, 2.5 KCl, 1.2 NaH_2_PO_4_, 30 NaHCO_3_, 20 HEPES, 25 glucose, 4 sodium ascorbate, 2 thiourea, 3 sodium pyruvate, 2 MgSO_4_(H_2_O)_7_, 2 CaCl_2_(H_2_O_2_), and 2 M NaOH added until pH reached 7.3-7.4 and osmolarity was 300-310 mOsm. Patch pipettes were pulled from 1.5 mm borosilicate glass capillaries (World Precision Instruments; 4-8 MW resistance). Pipettes were filled with K-gluconate intracellular solution for intrinsic experiments. K-gluconate composition was (in mM): 120 K-gluconate, 6 KCl, 10 HEPES, 4 ATP-Mg, 0.3 GTP-Na, 0.1 EGTA, and titrated to a pH of ∼7.2 with KOH. During recordings, slices were transferred to a perfusion chamber and continuously perfused with aCSF at 31 C at a rate of 4-7 mL/min: (in mM) 119 NaCl, 2.5 KCl, 1 NaH_2_PO_4_, 26 NaHCO_3_, 11 dextrose, 1.3 MgSO_4_(H_2_O)_7_, and 2.5 CaCl_2_(H_2_O_2_). All recordings were performed in the NAc core in mCherry positive cells that were visualized using CellSens software (Olympus). Cells were voltage-clamped at -70 mV and 15 current steps were injected starting at -100 pA and increasing in increments of 25 pA per step. For all patch-clamp experiments, cells collected from the same animal were grouped together for analysis. Elicited action potentials were recorded in current-clamp mode, counted, and analyzed using pClamp11 (Clampfit, Axon Instruments). Series access was monitored throughout recordings to ensure cells were only used if the access had not changed more than 10%. All drugs and reagents were obtained from Sigma-Aldrich.

#### Single-nuclei dissociation

Two weeks following surgery, animals (4 *lacZ*, 4 *Reln* 2M/2F per group) were briefly stunned prior to rapid decapitation. Brains were acutely blocked and NAc punches obtained, flash frozen, and stored at -80°C prior to nuclei isolation. Nuclei dissociation for snRNA-seq was performed as previously described^20^. One GEM (gel beads in emulsion) well was used per sample to generate barcoded nuclei (10× Genomics, PN-1000123, PN-1000122).

### Single-nucleus RNA-sequencing

#### Acute & Repeated Cocaine

Genes not expressed in at least 10 cells were removed before analysis on the integrated R object containing both acute and repeated cocaine and saline samples generated for Phillips et al^38^. Subclustering was performed by first subsetting all D1-MSNs from the integrated acute and repeated Seurat object^38^. Dimensionality reduction was performed on this subset of cells (DefaultAssay() = ‘Integrated’) via principal component analysis (PCA) retaining 17 PCs, followed by uniform manifold approximation and projection (UMAP) with 17 PCs as input. Cells were then clustered under a graph-based approach by first identifying K-nearest neighbors in PCA space, and then clustering under a Louvain algorithm with a resolution of 0.1. Gini coefficients were calculated using the gini() function that is available in the reldist v1.6-6 package^127,128^ using RNA expression values obtained with the Seurat function AverageExpression(). Differential gene expression testing was performed using DESeq2 v.1.40.2^122^ with a likelihood-ratio test to identify distinct genes among the D1-MSN clusters. Gini coefficient graphs were generated after merging the Gini coefficient matrix with the DESeq2 matrix for each cluster. Pearson correlations were calculated using the rmcorr v0.6.0 package with subject equal to the number of samples (n = 8) in the integrated dataset^129^.

#### Reln CRISPRi

Libraries were constructed according to manufacturer’s instructions using the Chromium Single Cell 3′ Library Construction Kit (10× Genomics, PN-1000190), which uses version 3 chemistry for gene expression. Nuclei (6,221) from adult rat NAc were sequenced on the Illumina NovaSeq6000 at the UAB Heflin Genomics core to a depth of ∼300,000 reads per nuclei. All raw fastq files were aligned to the Ensembl mRatBn7.2 (Rn7) genome using CellRanger v7.1.0^130^ with the associated Ensembl gene transfer format (gtf) file (version 105). CellRanger filtered outputs were analyzed with Seurat v4.3.0.1^131^ using R v4.3.1^132^ in RStudio v2023.06.1+524 as follows. First, to filter out background contamination, cell-free ambient RNA was removed using SoupX v.1.6.2^133^. Next, we continued quality filtering in R with Seurat v.4.3.0.1^131^ by removing nuclei containing <200 genes and >5% of reads mapping to the mitochondrial genome (5,846 nuclei post-processing). The counts data for each individual were then normalized, scaled by a factor of 10,000, and log-transformed. For each sample, we used Seurat to determine the 3,000 genes with highest cell-to-cell variability; these sets of genes were used to identify “anchor” pairs of cells between samples with a shared pattern of gene expression^134^. We then integrated the eight individual datasets based on the identified anchors, using the first 17 principal components (PCs). Downstream analysis was performed on the integrated dataset.

Expression for each gene in the integrated dataset was scaled to a mean of 0 and variance of 1 across cells. We used Seurat for initial dimensionality reduction and clustering as detailed below. We first performed dimensionality reduction via PCA retaining 17 PCs, followed by UMAP with the 17 previously identified PCs as input. Cells were then clustered under a graph-based approach by first identifying K-nearest neighbors in PCA space, and then clustering under a Louvain algorithm with a resolution of 0.3. After initial clustering but before differential gene expression analysis, we used DoubletFinder v.2.0.3^135^ to identify and remove heterotypic doublets, a consequence of two cells of different types being captured in a single droplet and thus presenting as a single heterotypic “cell.” We assumed a doublet formation rate of 4% based on 8,000 cells loaded per GEM well, as indicated in the 10x Genomics library preparation documentation. After removing doublets, we repeated the dimensionality reduction and clustering steps detailed above. We matched the resulting clusters to specific cell types based on known marker genes from previous work^38^.

To identify genes differentially expressed between *Reln* knockdown and *lacZ* controls, we conducted a differential expression analysis on a pseudobulked gene expression matrix created by summing counts for each gene within a sample (or single GEM well) and across cells within each cell type. Genes with mean counts <5 were removed. Differential expression analysis was conducted with DESeq2 v.1.40.2^122^ using a likelihood ratio test (LRT) with treatment condition as the main effect and controlling for the effect of sex to ensure that identified significant differential expression could be attributed to the treatment and not confounded by differences in response between sexes. To assess differential expression among cell types, we conducted a separate LRT with cell type as the main effect and controlling for sex. All R code for our snRNA-seq analyses is available at https://github.com/Jeremy-Day-Lab/Brida_Reln_2024.

To correlate cell types between this study and prior work ^38^, the function FindMarkers() in the Seurat package was used to define marker genes using the default Wilcoxon Rank Sum test for both datasets. Marker gene sets were filtered for genes with a pAdj < 0.05. Marker gene lists and the relative enrichment (log_2_(fold-change)) relative to all other clusters for each cell type were then concatenated, with annotation as to which dataset each cell type originated from. Pearson’s correlation was then calculated using the cor() function in R.

Gene ontology analysis was performed using the online tool, g:Profiler ^136^ using DEGs defined by DESeq2 with pAdj < 0.05 in the D1-MSN cluster (28 genes; **Table S1**). A custom gene set was used as background reference, consisting of all genes detected in the D1-MSN cluster (21,042 genes), with the “custom over all” option selected and Bonferroni was used for statistical correction.

Pseudotime calculations were performed using Monocle v3_1.3.5^137^ and Seurat v4.3.0^131^. A Seurat object containing D1-MSNs from male and female adult Sprague-Dawley rats from *lacZ* control and *Reln* knockdown groups (2M/2F per group) was subject to the standard dimensionality reduction and clustering workflow using 17 principal components and a resolution value of 0.1. *Reln* expression was removed from the RNA matrix prior to subclustering. Annotation, counts metadata, and cell barcode information were extracted from the Seurat object and reconstructed into a Monocle v3 cds object using new_ cell_data_set(). Rather than reclustering, the UMAP coordinates were extracted from the Seurat object and added to the Monocle object. The trajectory graph was created using learn_graph() and the root cell was chosen in the ‘inactivated’ population that expressed *Drd1* and *Htr4*^*38*^. Pseudotime values for each cell were exported from the cds object and added back to the Seurat object metadata.

### Statistical Analysis

Sample sizes were determined using a freely available calculator (https://www.stat.ubc.ca/~rollin/stats/ssize/n2.html). Transcriptional differences from RT-qPCR experiments were compared with an unpaired *t*-test with Welch’s correction, one-way ANOVA with Tukey’s *post hoc* tests, or two-way ANOVA with Sidak’s *post hoc* tests, where appropriate. CPP data were compared with a two-way ANOVA with Tukey’s multiple comparisons test. smRNA-FISH expression differences were compared with nested t-test analysis with correction for multiple comparisons where appropriate. Statistical significance was designated at α = 0.05 for all analyses. Statistical and graphical analyses were performed with Prism software (GraphPad, La Jolla, CA). Statistical assumptions (e.g. normality and homogeneity for parametric tests) were formally tested and examined via boxplots. Full details on all statistical tests can be found in **Table S3**.

## DATA AVAILABILITY

All relevant data that support the findings of this study are available by request from the corresponding author (J.J.D.). Custom code can be found at https://github.com/Jeremy-Day-Lab/Brida_Reln_2024. Sequencing data that support the findings of this study are available in Gene Expression Omnibus. Accession numbers of specific datasets are outlined below. Bulk RNA-seq *Reln* CRISPRi: GSE269366 snRNA-seq *Reln* CRISPRi: GSE269490

## ACKNOWLEDGEMENTS

We thank all current and former Day Lab members for assistance and support, as well as Natalie Davis, Katherine Savell, and Nicholas Boyle for assistance. This work was supported by NIH grants DP1DA039650 & R01DA053743 (JJD), F30DA057821 (KLB), R01DA053581 (KM), the McKnight Foundation Neurobiology of Brain Disorders Award (JJD) and the Lieber Institute for Brain Development (KM, KRM). LI is supported by the Civitan International Research Center at UAB. KLB also received support from the Civitan International Research Center Emerging Scholars Award. We acknowledge support from the University of Alabama at Birmingham Biological Data Science Core (RRID:SCR_021766), the UAB Heflin Center for Genomic Sciences, and the UAB Flow Cytometry & Single Cell Core Facility.

## AUTHOR CONTRIBUTIONS

KLB and JJD conceived of rat model experiments, which were completed by KLB with assistance from ETJ, RAP, JJT, EKM, and MEZ. KLB, JJD, CEN, LI, and RAP performed bioinformatics analysis. JJD supervised all work at the University of Alabama at Birmingham. KDM, MT, TMH, KRM, and KM designed and completed human RNAscope experiments, which were supervised by KM at the Lieber Institute for Brain Development. KLB and JJD wrote the manuscript with assistance from all authors. All authors have approved the final version of the manuscript.

---

**Figure S1.**
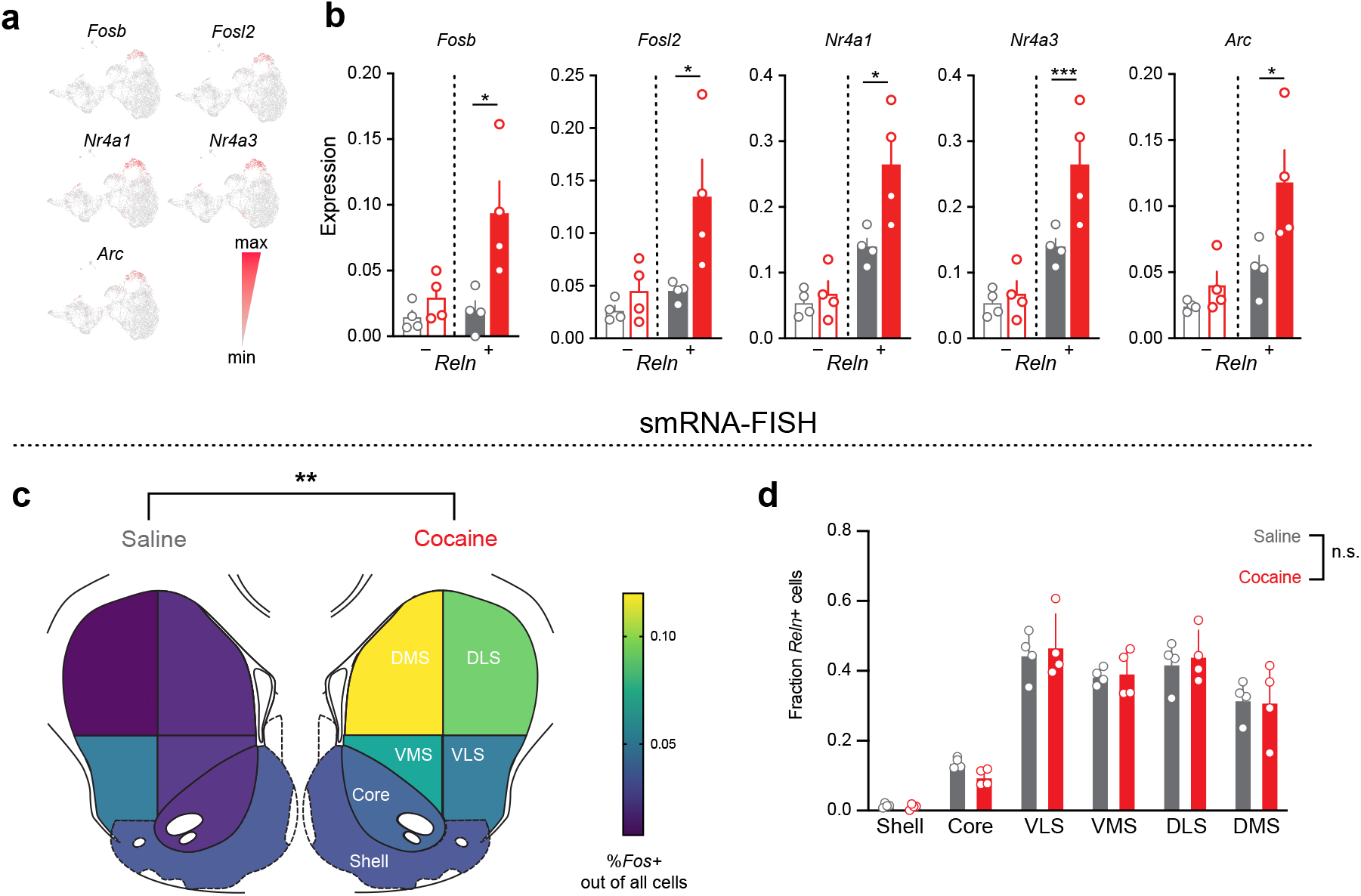
*Reln* serves as a marker of Drd1-MSNs with a robust transcriptional response to cocaine but does not change with cocaine exposure. **a**, Global UMAP showing 16 transcriptionally distinct cell populations within the NAc. Inset shows subclustering of the Drd1-MSN population into active (red) and inactive (gray; i1-4) groups. **b**, Drd1-MSN subcluster feature plots showing expression of select immediate early genes, from left ot right: *Fosb, Fosl2, Nr4a1, Nr4a3*, and *Arc*. **c**, snRNA-seq expression data for select immediate early genes across all Drd1-MSNs, split by *Reln* expression, from left ot right: *Fosb, Fosl2, Nr4a1, Nr4a3*, and *Arc* (nested t-test; \**p*<0.05, \*\*\**p*<0.001). **c**, Heatmap of showing percent out of total cells that are *Fos+* in Saline (left) and Cocaine (right) animals (unpaired t-test of Saline vs Cocaine, *p* < 0.01). **d**, *Reln* expression Drd1-MSNs in the nucleus accumbens does not change with cocaine (*p* > 0.05, nested t-test).

**Figure S2.**
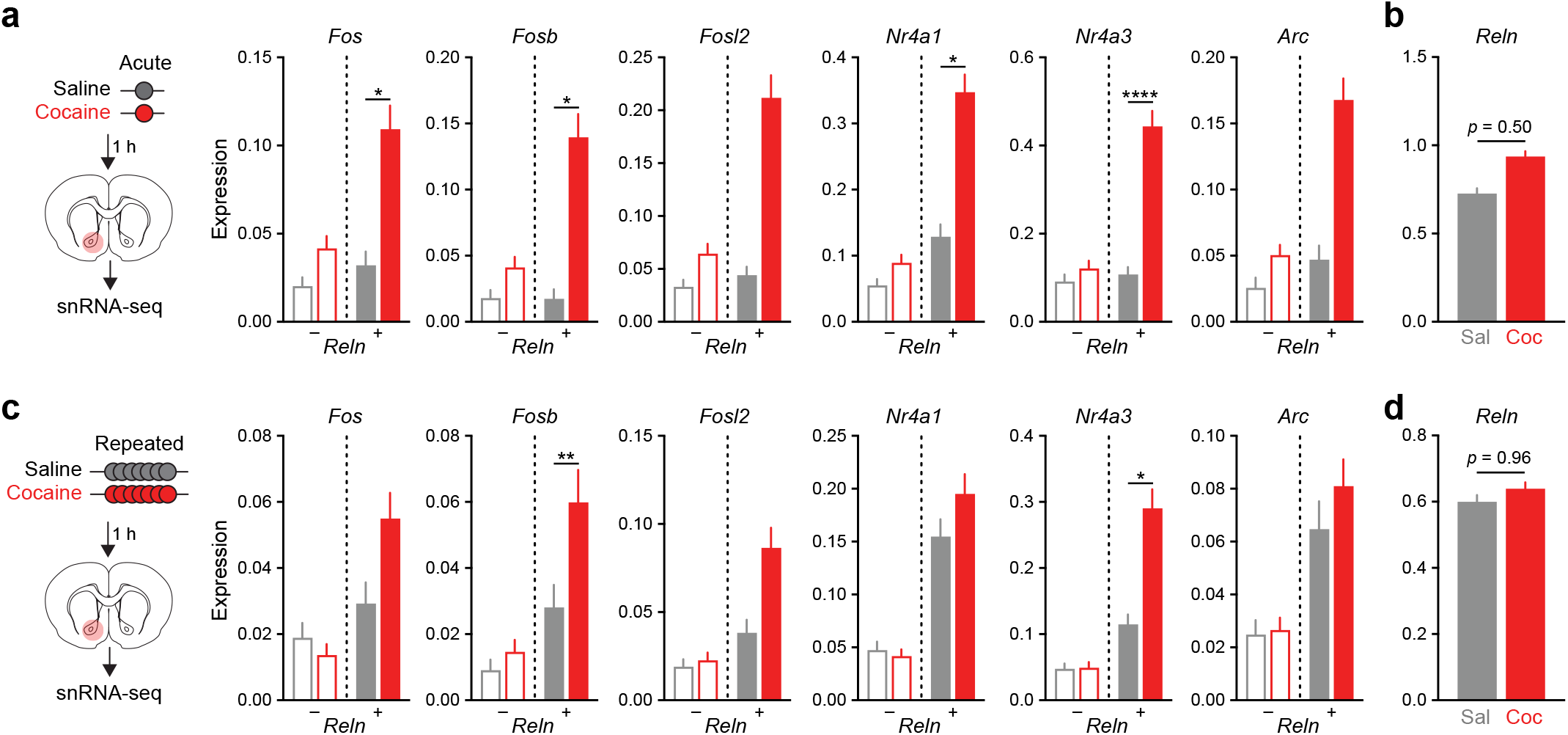
*Reln* marks cocaine-sensitive Drd1-MSNs following acute or repeated cocaine. **a**, Immediate early genes increase in *Reln*+ Drd1-MSNs but not *Reln-*Drd1-MSNs following acute cocaine. **b**, *Reln* mRNA levels do not change following acute cocaine (nested t-test, *p* = 0.55). **c**, Immediate early gene expression is induced in *Reln*+ Drd1-MSNs following repeated cocaine. **d**, *Reln* mRNA levels do not change following repeat-ed cocaine (nested t-test, *p* = 0.96).

**Figure S3.**
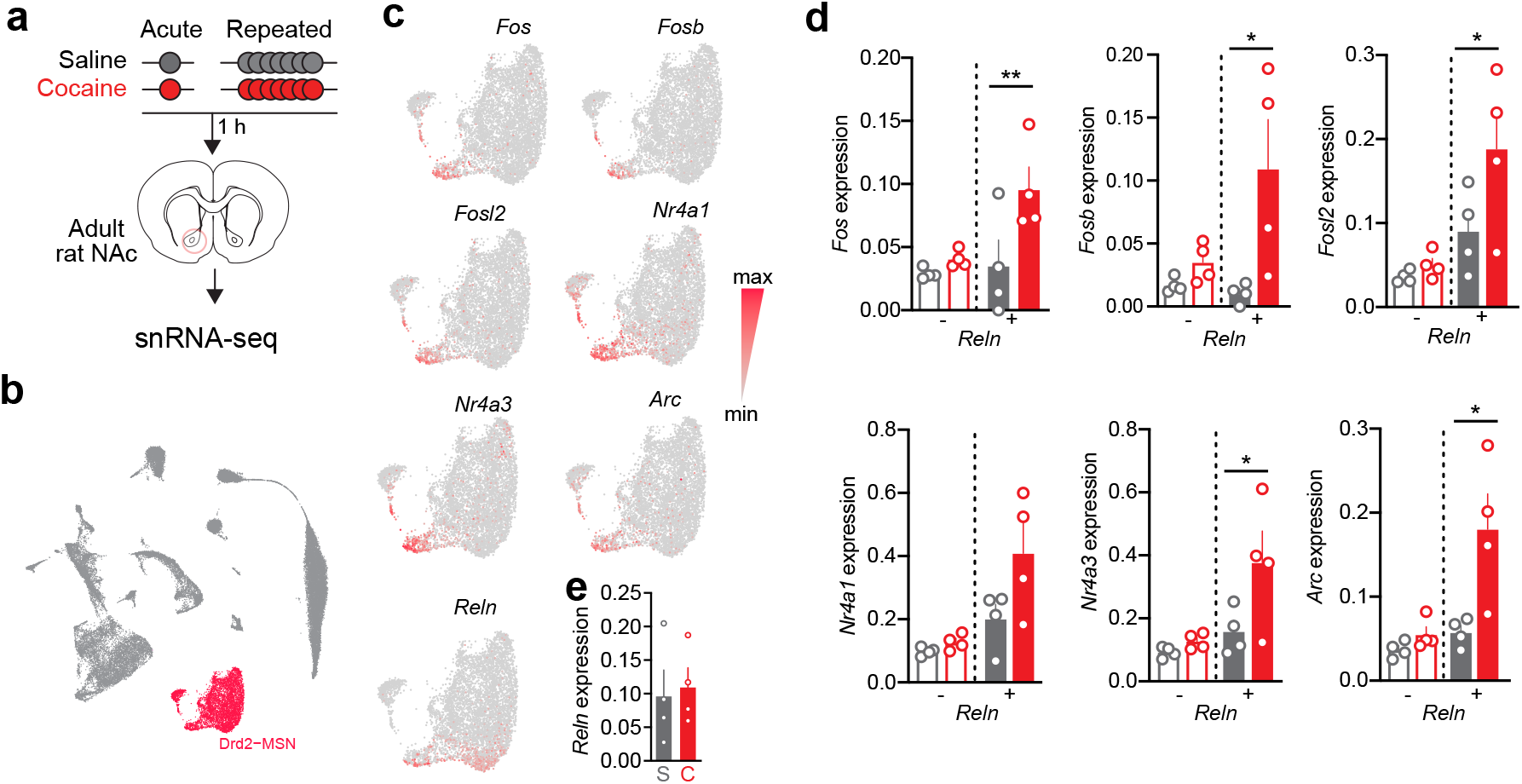
*Reln* serves as a marker of Drd2-MSNs with a robust transcriptional response to cocaine but does not change with cocaine exposure. **a**, snRNA-seq workflow for acute and repeated saline (gray/S) or cocaine (red/C). **b**, Global UMAP showing 16 transcriptionally distinct cell populations within the NAc, Drd2-MSN populations highlighted in red. **c**, Drd2-MSN cluster feature plots showing expression of select immediate early genes, from left ot right: *Fos, Fosb, Fosl2, Nr4a1, Nr4a3*, and *Arc*. **d**, snRNA-seq expression data for select immediate early genes across all Drd2-MSNs, split by *Reln* expression, from left ot right: *Fos, Fosb, Fosl2, Nr4a1, Nr4a3, Arc*, and *Reln* (nested one-way ANOVA; \**p*<0.05, \*\**p*<0.01). **e**, *Reln* expression Drd2-MSNs in the nucleus accumbens does not change with cocaine (*p* = 0.79, nested t-test).

**Figure S4.**
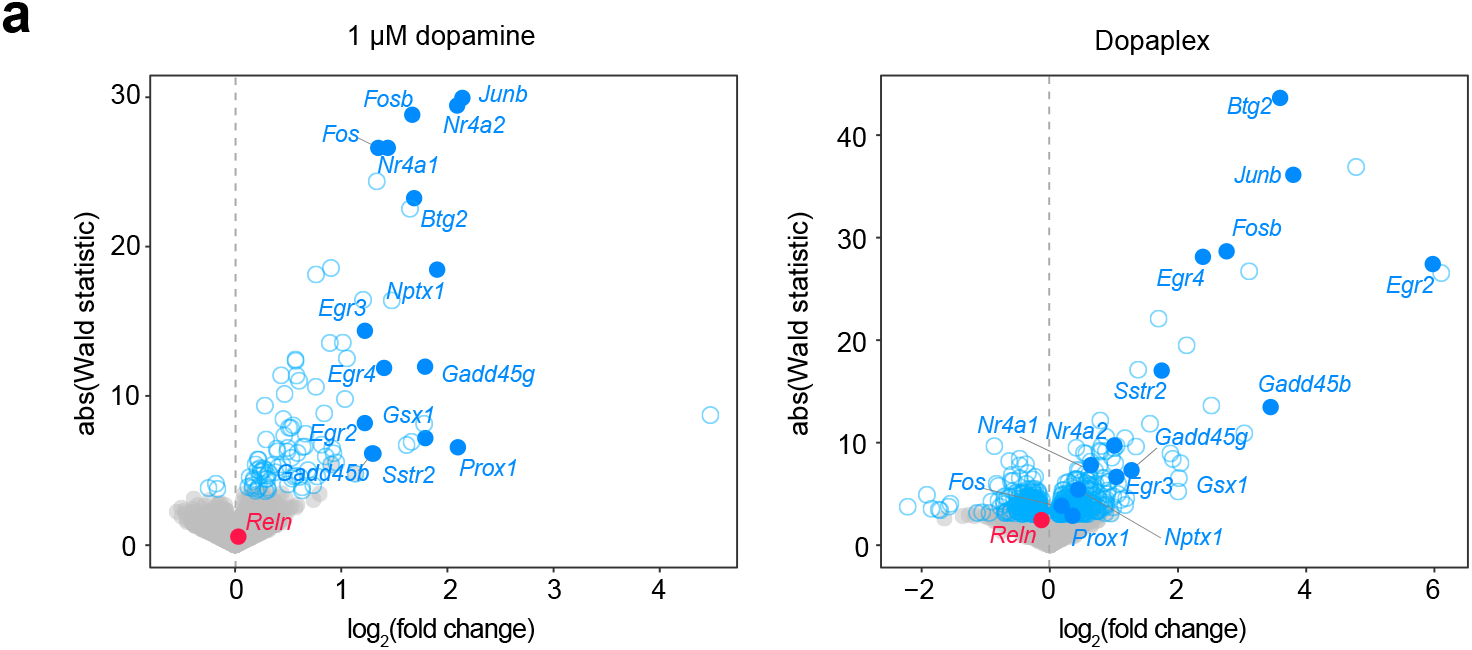
*Reln* expression is unchanged following either dopamine or Dopaplex treatment. **a**, Volcano plot of dopamine-induced DEGs (left; *Reln p* adjusted > 0.99) and Dopaplex-induced DEGs (right; *Reln p* adjusted = 0.15). Dopaplex targets highlighted in blue, *Reln* highlighted in red, grey indicates non-siginficant genes.

**Figure S5.**
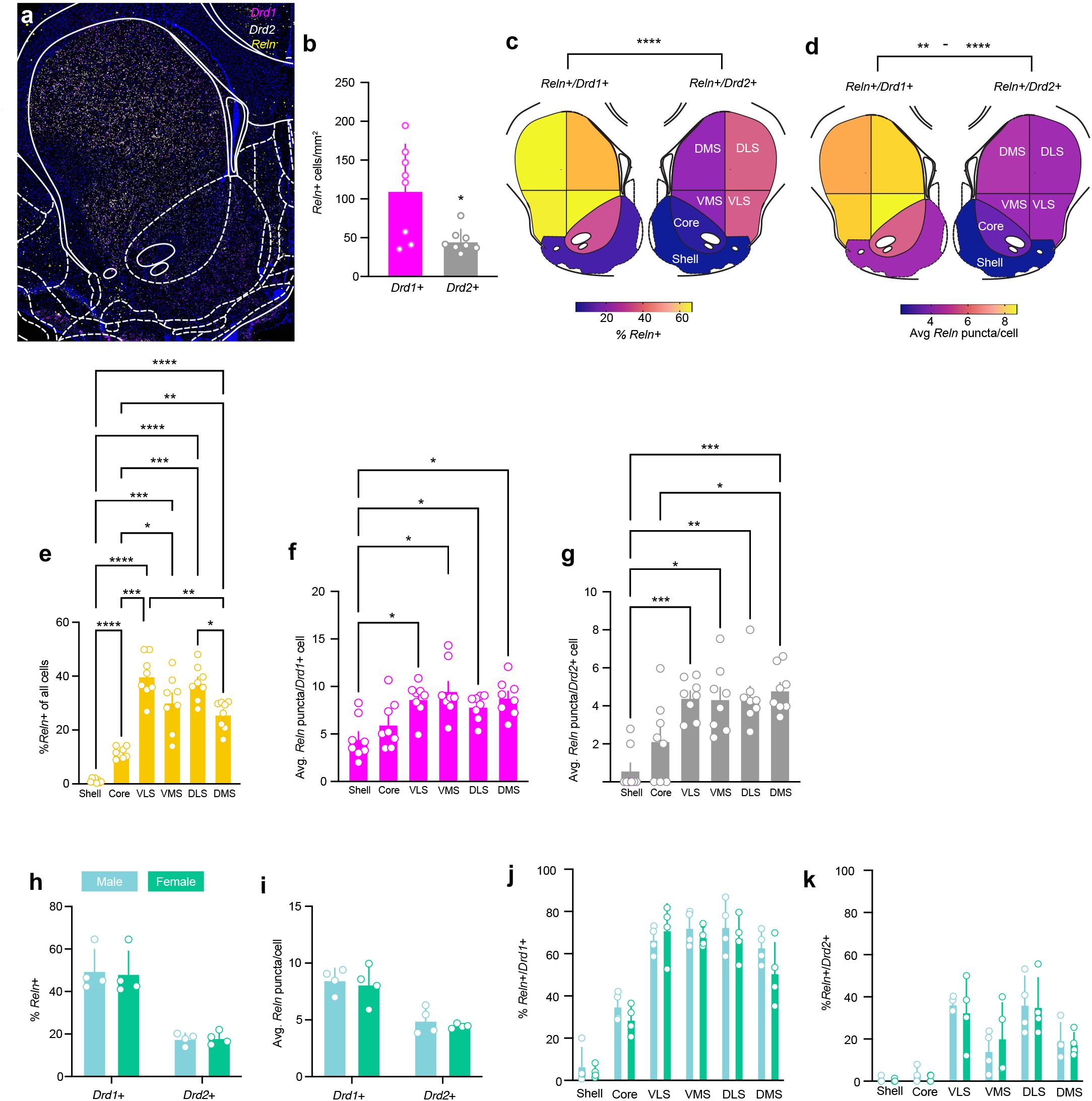
RNAscope shows Drd1-MSN enrichment in *Reln*. **a**, RNAscope image of rat striatum with *Reln* (yellow), *Drd1* (magenta), and *Drd2* (white). Scale bar 500 µm. **b**, Density of *Reln*+ cells throughout the striatum (*p*<0.05, paired t-test). **c**, Heatmap of percent *Reln*+ *Drd1* (left) or *Drd2* (right) by striatal subregion (nucleus accumbens core (Core), nucleus accumbens shell (Shell), ventromedial striatum (VMS), ventrolateral striatum (VLS), dorsomedial striatum (DMS), dorsolateral striatum (DLS); two-way ANOVA with repeated measures (regional differences main effect F (5, 35) = 65.77, *p* < 0.0001; *Drd1* vs *Drd2* main effect F (1, 7) = 515.5, *p* < 0.0001; interaction, F (5, 35) = 18.71, *p* < 0.0001); Tukey’s post-hoc mutliple comparsons, *Drd1* vs *Drd2* shell *p* > 0.05; *Drd1* vs *Drd2* all other regions *p* < 0.0001). **d**, Heatmap of average *Reln* puncta per cell by striatal subregion (NAc core, NAc shell, VLS, VMS, DLS, & DMS; two-way ANOVA with repeated measures (regional differences main effect, F (5, 35) = 17.84, *p* < 0.0001; *Drd1* vs *Drd2* main effect F (1, 7) = 99.01, *p* < 0.0001; interaction, F (5, 35) = 0.5438, *p* > 0.05); Tukey’s post-hoc comparson *Drd1* vs *Drd2*: shell, core, DLS, & DMS *p* < 0.01; VMS *p* < 0.0001; VLS *p* < 0.001)). **e**, Percent *Reln*+ cells by region (\**p* < 0.05, **p < 0.01, \*\*\**p* < 0.001, \*\*\*\**p* < 0.0001, one-way ANOVA). **f**, Average *Reln* puncta in *Drd1*+ cells by region (\**p* < 0.05, \*\**p* < 0.01, \*\*\**p* < 0.001, one-way ANOVA). **g**, Average *Reln* puncta in *Drd2*+ cells by region (\**p* < 0.05, \*\**p* < 0.01, \*\*\**p* < 0.001, one-way ANOVA). **h**, Comparison of *Reln* positivity in *Drd1*+ and *Drd2*+ between sexes shows no difference (*p* > 0.05, 2-way ANOVA). **i**, Comparison of *Reln* expression levels in *Drd1*+ and *Drd2*+ between sexes shows no difference (*p* > 0.05, 2-way ANOVA). **j**, *Reln* expression in *Drd1*+ cells across striatal subregions is not different between sexes (*p*>0.05, 2-way ANOVA with Tukey’s post-hoc correction). **k**, *Reln* expression in *Drd2*+ cells across striatal subregions is not different between sexes (*p*>0.05, 2-way ANOVA with Tukey’s post-hoc correction).

**Figure S6.**
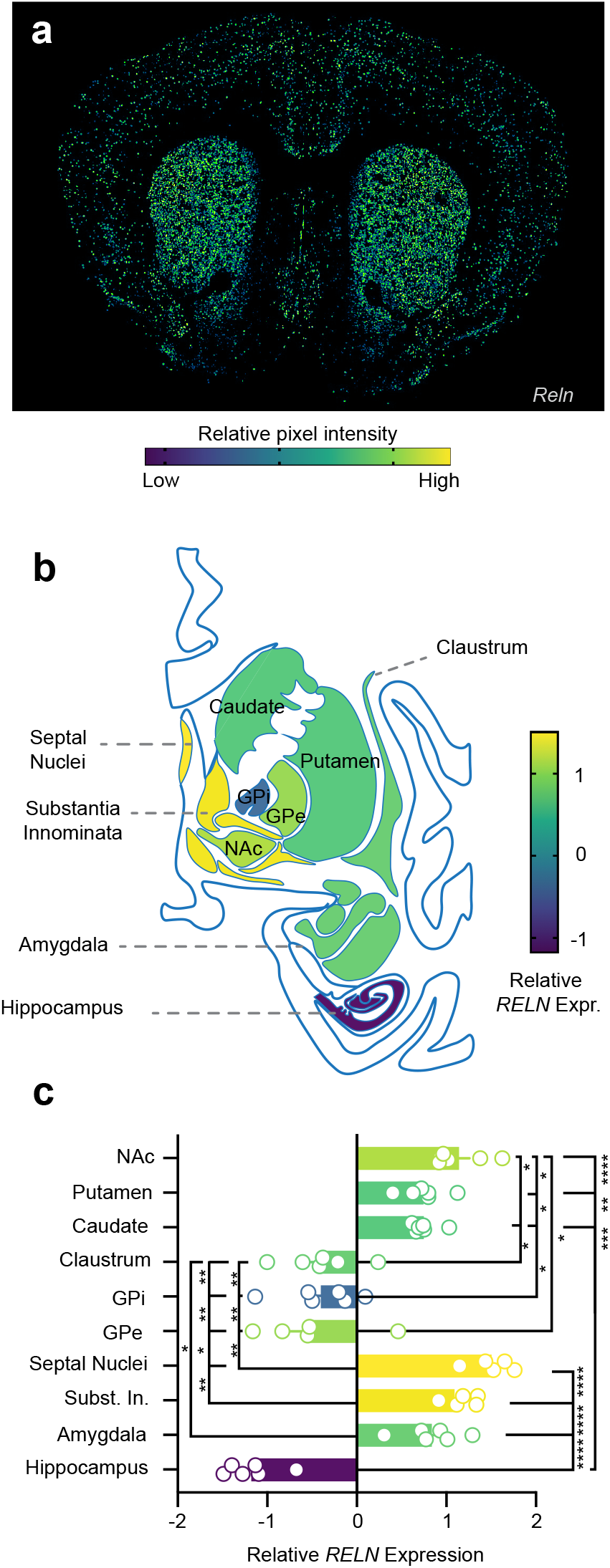
*Reln* expression in mouse and human striatum. **a**, *Reln* expression in mouse striatum (https://mouse.brain-map.org/gene/show/19462), colored based on pixel intensity. **b**, Heatmap of *RELN* expression in human striatum, averaged across 6 human donors (NAc: Nucleus Accumbens; GPi: Globus pallidus interanl; GPe: Globus pallidus external. All data from the Allen Brain Atlas) demonstrates striatal enrichment of *RELN*. **c**, Quantification of *RELN* expression in B (1-way ANOVA with Tukey’s correction for mutliple comparisons; * *p* < 0.05, \*\**p* < 0.01, **** *p* < 0.0001).

**Figure S7.**
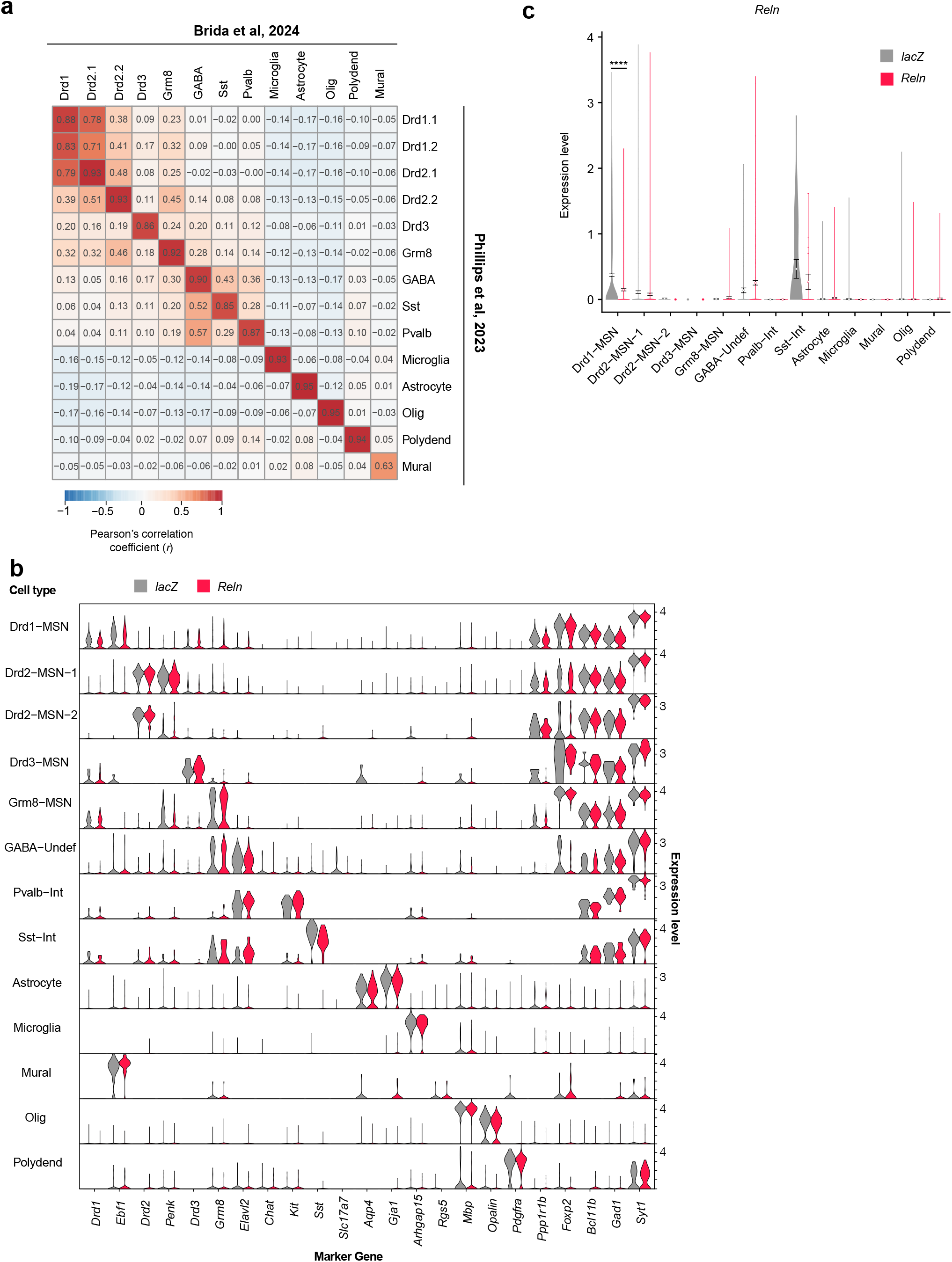
Validation of snRNA-sequencing dataset. **a**, Correlation matrix showing high agreement of cell identities between this study and Phillips et al, 2023. **b**, Marker gene expression by cell type split by gRNA target. **c**, *Reln* expression across cell types, split by gRNA target (LRT, **** = p.adj < 0.0001).

**Figure S8.**
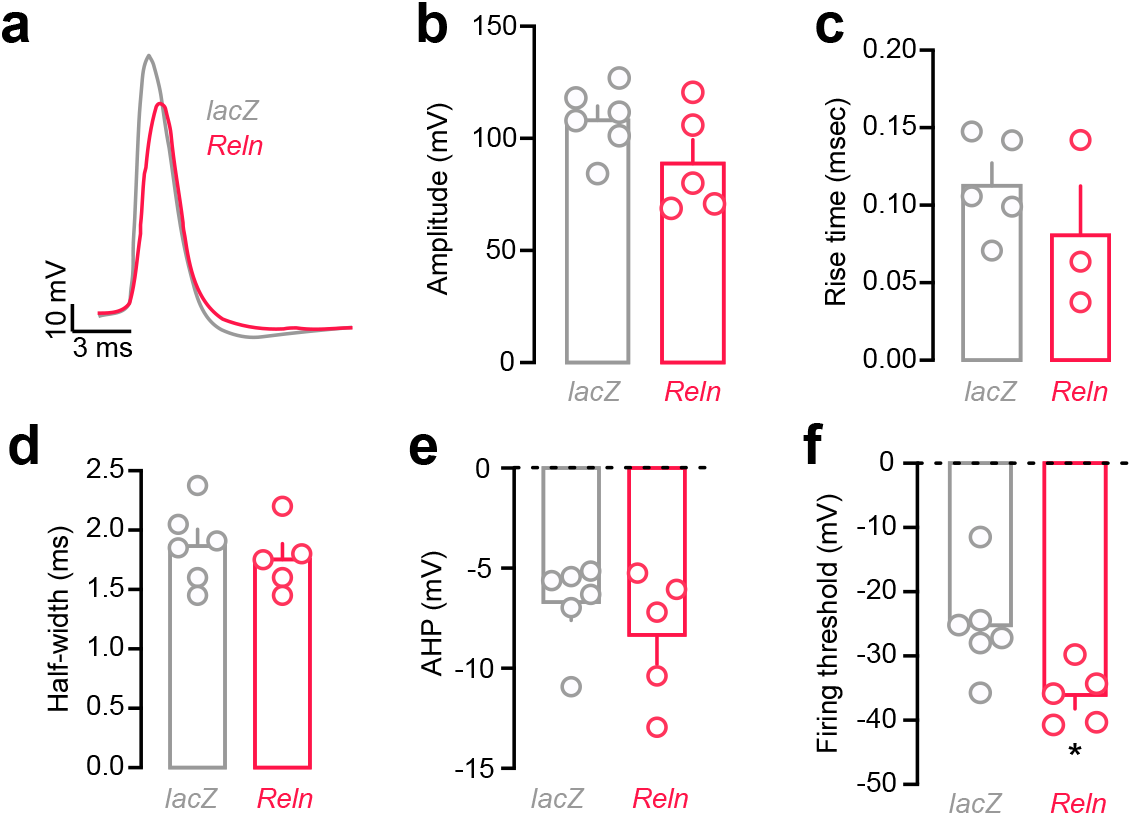
CRISPR-mediated *Reln* knockdown does not impact action potential forms but decreases firing threshold. **a**, Represne-stative action potential (AP) traces from *lacZ* gRNA (gray) or *Reln* gRNA (red) obtained during current injection. **b-e**, *Reln* does not affect AP amplitude (**b**), rise time (**c**), half-width (**d**), or after-hyper polarization (AHP (**E**); *lacZ* n = 6 animals, 17 cells; *Reln* n = 5 animals, 11 cells). **f**, *Reln* cells demonstrate lower firing threshold (*Reln* avg. threshold = -36.2 mV, *lacZ* avg. threshold = -25.3 mV; *p*=0.0122, nested t-test).

**Figure S9.**
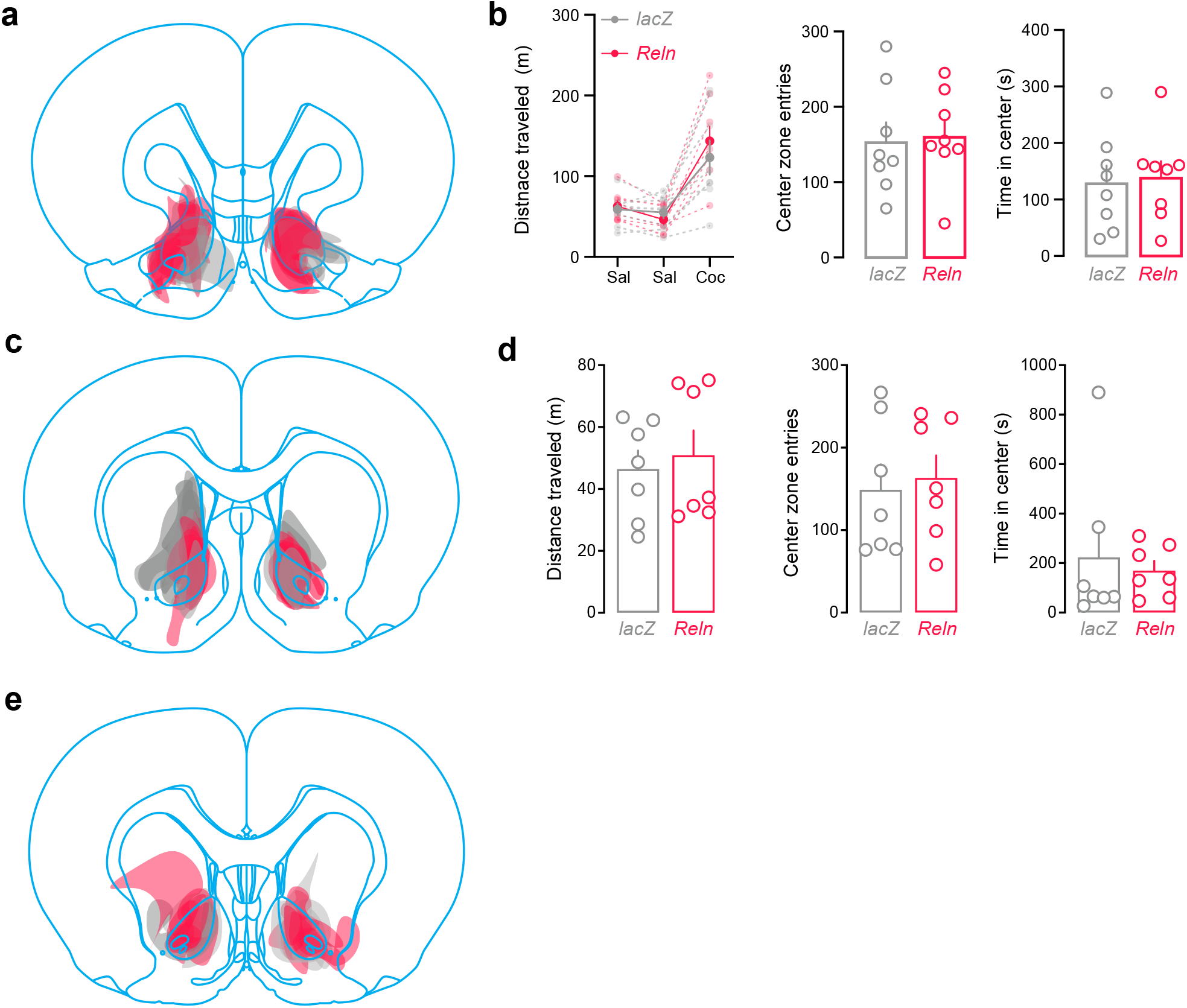
Viral targeting and baseline locomotion for behavioral studies. **a**, Virus targeting for locomotor sensitization (n = *lacZ* (gray), *Reln* (red)). Atlas approximately +2.76 mm from Bregma (Watson & Paxinos). **b**, Locomotion across saline Day 1 & 2 and cocaine Day 3 did not differ between *lacZ* and *Reln* gRNA groups (2way ANOVA main effect of treatment p < 0.001, main effect gRNA p = 0.72, interaction p = 0.31; Sudak multiple comparisons *lacZ* vs *Reln* gRNA: Saline 1 *p* > 0.99, Saline 2 *p* = 0.95, Cocaine 1 *p* = 0.60). **c**, Center zone entries and time in center did not differ between gRNA groups (t-test, entries *p* = 0.83, time *p* = 0.81; data from saline Day 1). **d**, Virus targeting for CPP (n = 14 *lacZ* (gray), 11 *Reln* (red)). Atlas approximately +2.04 mm from Bregma (Watson & Paxinos). **e**, Baseline locmotion, time in center, and center zone entries did not differ between *lacZ* and *Reln* gRNA groups. **f**, Virus targeting for IVSA (n = 6 *lacZ* (gray), 8 *Reln* (red)). Atlas approximately +2.28 mm from Bregma (Watson & Paxinos).

